# Magnitude and kinetics of the human immune cell response associated with severe dengue progression by single-cell proteomics

**DOI:** 10.1101/2022.09.21.508901

**Authors:** Makeda L. Robinson, David R. Glass, Veronica Duran, Olga Lucia Agudelo Rojas, Ana Maria Sanz, Monika Consuegra, Malaya Kumar Sahoo, Felix J. Hartmann, Marc Bosse, Rosa Margarita Gelvez, Nathalia Bueno, Benjamin A. Pinsky, Jose G. Montoya, Holden Maecker, Maria Isabel Estupiñan Cardenas, Luis Angel Villar Centeno, Elsa Marina Rojas Garrido, Fernando Rosso, Sean C. Bendall, Shirit Einav

## Abstract

Approximately five million dengue virus-infected patients, particularly children, progress to a potentially life-threatening severe dengue (SD) infection annually. To identify the immune features and temporal dynamics underlying SD progression, we performed deep immune profiling by mass cytometry of PBMCs collected longitudinally from SD progressors (SDp) and uncomplicated dengue (D) patients. While D is characterized by early activation of innate immune responses, in SDp there is rapid expansion and activation of IgG-secreting plasma cells and memory and regulatory T cells. Concurrently, SDp, particularly children, demonstrate increased proinflammatory NK cells, inadequate expansion of CD16^+^ monocytes, and high expression of the FcγR, CD64 on myeloid cells, yet diminished antigen presentation. Syndrome-specific determinants include suppressed dendritic cell abundance in shock/hemorrhage vs. enriched plasma cell expansion in organ impairment. This study reveals uncoordinated immune responses in SDp and provides insights into SD pathogenesis in humans with potential implications for prediction and treatment.

## Introduction

Dengue virus (DENV) is a major threat to global health, infecting approximately 400 million people annually in over 130 countries (Bhatt et al., 2013; WHO, 2012). Each year, 3–6 million symptomatic dengue patients develop severe dengue (SD) within days, which can be life-threatening (Stanaway et al., 2016; WHO, 2012; Zeng et al., 2021). To identify patients at risk for SD progression, the World Health Organization (WHO) defined clinical criteria that distinguish dengue patients with warning signs (DWS) from those with uncomplicated dengue (D) and recommended their close monitoring (WHO, 2012). Nevertheless, since warning signs for SD often develop late in the course of illness and are nonspecific, their implementation does not capture all SD progressors (SDp) and has increased hospital care burden (Hadinegoro, 2012; Narvaez et al., 2011). While we and others have recently identified candidate clinically usable biomarkers for early identification of SDp (Liu et al., 2022; Robinson et al., 2019; Tissera et al., 2017; Zanini et al., 2018), additional validation is ongoing and novel biomarkers are needed.

The best characterized risk factor for SD is secondary infection with a heterologous DENV serotype causing antibody-dependent enhancement (ADE) of infection (Halstead and O’Rourke, 1977; Halstead and O’rourke, 1977; Katzelnick et al., 2017). Additional factors including increased plasma cells (Kwissa et al., 2014; Nascimento et al., 2014), aberrant T cell responses (Mongkolsapaya et al., 2006), cytokine storm (Chen et al., 2008; Mongkolsapaya et al., 2006), and high and prolonged viremia have been implicated in the development of SD (Wang et al., 2006). Yet, since our understanding of SD pathogenesis is largely based on cultured cells and immunocompromised mouse studies, the role of these and other factors in natural infection in humans is poorly characterized. Moreover, it remains unknown why DENV-infected children are at greater risk for SD (Guzmán et al., 2002; Wichmann et al., 2004), and why SD manifests as dengue hemorrhagic fever (DHF) and dengue shock syndrome (DSS) in some but as organ impairment (OI) in others (WHO, 2012).

To address these gaps in knowledge, we applied mass cytometry (CyTOF), a proteomic single-cell approach, to 124 peripheral blood mononuclear cell (PBMC) samples from our dengue Colombia cohort. The unique design and clinical heterogeneity of our cohort facilitated comprehensive analysis of immune cell subtype abundance and functional marker expression revealing distinct and overlapping responses between: (1) SDp and non-progressors; (2) SDp adults and children; and (3) SDp who manifested with DHF/DSS or OI. Longitudinal sample analysis profiled the kinetics of the immune response starting prior to progression to SD through convalescence, revealing dysregulation of the temporal switch between innate and adaptive immune responses associated with early exuberant immune regulation in SDp.

## Results

### High-dimensional immune profiling delineates disease severity in dengue infection

We collected longitudinal PBMC and serum samples from adults and children enrolled in the dengue Colombia cohort during acute infection and at convalescence. We performed CyTOF analysis on a total of 124 samples from healthy controls C (n=15), D (n=40), DWS (n=23), and SDp (n=22) patients of whom 19 progressed to SD within several days following enrollment and 3 met criteria of SD at presentation (**Figure 1A, Table S1.1, Table S1.2, Table S1.3**). The majority of patients were previously exposed to DENV (see methods), as expected for an endemic region (**Figure S1A**). To broadly profile the immune response, we employed an antibody panel targeting 36 cellular markers encompassing proteins associated with immune activation, immune checkpoint, and Fc-binding.

Following singlet gating, normalization, and quality filtering (**Figures S1B-D**), the full dataset consisted of 9,051,718 cells. To capture patterns of cell abundance and protein expression at different levels of granularity, we clustered cells at three resolutions: 4 cell types (B, NK, myeloid, and T cells) (**Figure 1B**, panels), 11 cell subtypes (**Figure 1C**, colors), and 29 cell populations (**Figure 1B**, individual rows). We used expression of lineage-specific proteins (**Figure 1B**, columns) to cluster cells into predefined immune cell subtypes so that differential expression of activation, checkpoint, and Fc-binding proteins would not bias clustering (see methods). Differential abundance of immune cell phenotypes was evident by UMAP analysis (Becht et al., 2019) of the entire cohort (**Figure 1C**) and the various disease severity categories (**Figure S1E**), regardless of cluster labels.

**Figure 1.**
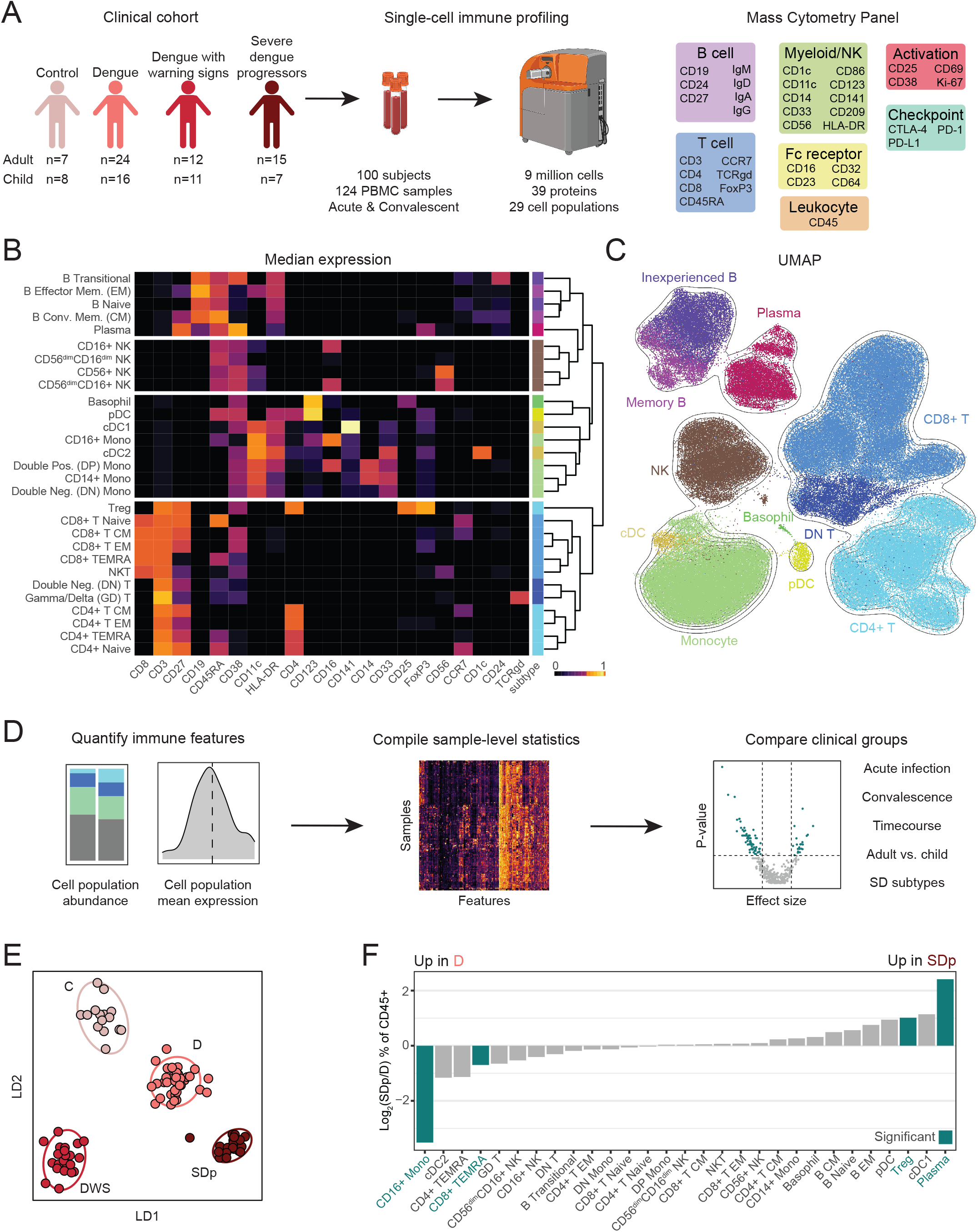
High-dimensional immune profiling delineates disease severity in dengue infection. (A) Experimental approach. Acute and convalescent PBMC samples were collected from a clinical cohort of DENV-infected patients and healthy controls. Samples were analyzed by mass cytometry using an antibody panel targeting proteins expressed by multiple immune cells. (B) A heat map of the median scaled arcsinh-transformed expression (color coded) of lineage molecules (columns) in cell B, NK, myeloid, and T cells populations (rows) spaced by cell type. Cell subtype color annotation is depicted on the right. PBMCs were equally subsampled by clinical status and patient. (C) UMAP generated using all molecules as input colored by cell subtype (B). PBMCs were equally subsampled by clinical status and patient. (D) Analysis approach. Immune features were quantified and compiled into summary statistics for each sample. Distribution tests were performed for each feature for various comparisons. (E) Linear discriminate analysis separating acute patient samples by clinical status. Dots represent individual patients and ellipses represent 95% confidence intervals (CI). The manifold was derived using all features significantly different in pairwise comparisons of clinical statuses. F) Log_2_ ratio of median abundances of cell populations (columns) between D and SDp patients out of total CD45+ cells. Teal bars indicate significance (q<0.05 & |effect|>0.5) via Wilcoxon rank sum tests. Q-values represent FDR-corrected p-values.

To quantitatively capture the state of the immune system, we compiled a set of 378 features consisting of cell population abundances and mean expression values for each sample (**Figure 1D**). With this feature set, we performed pairwise distribution tests to identify features significantly different between severity categories in the setting of acute infection (within 12 days of fever onset) (**Table S2**), convalescence **(Table S3**), adults and children **(Table S4)**, SD categories **(Table S5) (Figure 1D),** and early acute timepoints (within 8 days of infection) (**Table S6**).

To compare immune features between disease severity categories, we compiled samples collected within 12 days after reported symptom onset. When several samples were available, we used only the earliest sample to enrich for acute infection and avoid bias from over-representation of any single patient. For all features, we performed pairwise Wilcoxon rank sum tests between disease categories, corrected for multiple hypotheses (false-discovery rate (FDR) method), and quantified effect sizes (**Table S2**). Linear discriminant analysis (LDA) (Amouzgar et al., 2022) using all features that were significantly different between any two severity categories (q<0.05 & |effect size| >0.5) effectively separated the dengue severity categories (**Figure 1E**), confirming that our analytic framework captured substantial differences in the immune state.

To visualize differences in cell population abundances between SDp and D patients, we plotted the log_2_ SDp/D ratio of median abundances (% of CD45^+^ cells) (**Figure 1F**). The abundance of plasma cells (q=0.003) and regulatory T cells (Tregs) (q=0.025) was significantly greater in SDp than D patients. In contrast, the abundance of CD16^+^ monocytes (q=0.002) and CD8^+^ T effector memory cells re-expressing CD45RA (EMRA) associated with cellular senescence and terminal differentiation (Callender et al., 2018), was significantly lower in SDp than D (q=0.018). Trends towards expansion of conventional type 1 dendritic cells (cDC1s; stimulate CD8^+^ T cell responses by cross-presentation (Collin and Bigley, 2018)) and plasmacytoid dendritic cells (pDCs; type I IFN producing (Gilliet et al., 2008)), concurrent with a reduction of cDC2s (predominately prime CD4^+^ T cells (Krishnaswamy et al., 2017) and CD4^+^ TEMRAs in SDp relative to D were also observed. Altogether, these results demonstrate that our experimental and analytical approaches capture differences in immune cell subtype abundance with differential disease severity during the acute phase of DENV infection.

### Immune activation and regulation are simultaneously enriched in SDp

Since plasma cells were more abundant in SDp patients, we compared their abundance across all disease severity categories. While lower than SDp, D patients had significantly higher plasma cell abundance than C (q=0.027), as expected during an acute viral infection (**Figure 2A**). Nonetheless, SDp had higher plasma cell abundance (median=7.64%) than DWS (median=1.85%, q=0.044), D (median=1.44%, q=0.003), and C subjects (median=0.40%, q= 0.00006). Class-switched plasma cells in SDp were enriched for IgG over IgA (q=0.040) (**Figure 2B**) and expressed greater levels of Ki-67 relative to D (q=0.005) (**Figure 2C**), indicating greater production of the antibody isotype associated with ADE (Bournazos et al., 2020) and cell proliferation, respectively. Plasma cell abundance and plasma cell IgG usage diminished in both D and SDp in convalescent samples (**Fig. S2A-B**). Mean Ki-67 expression in plasma cells in SDp remained high in convalescence and in D, was significantly increased in convalescent samples over acute samples (p=0.0002) (**Fig. 2D**). These convalescent expression patterns contrast expression in C (**Fig. 2C**), suggesting that DENV infection induced heightened plasma cell activation even weeks to months after acute infection.

**Figure 2.**
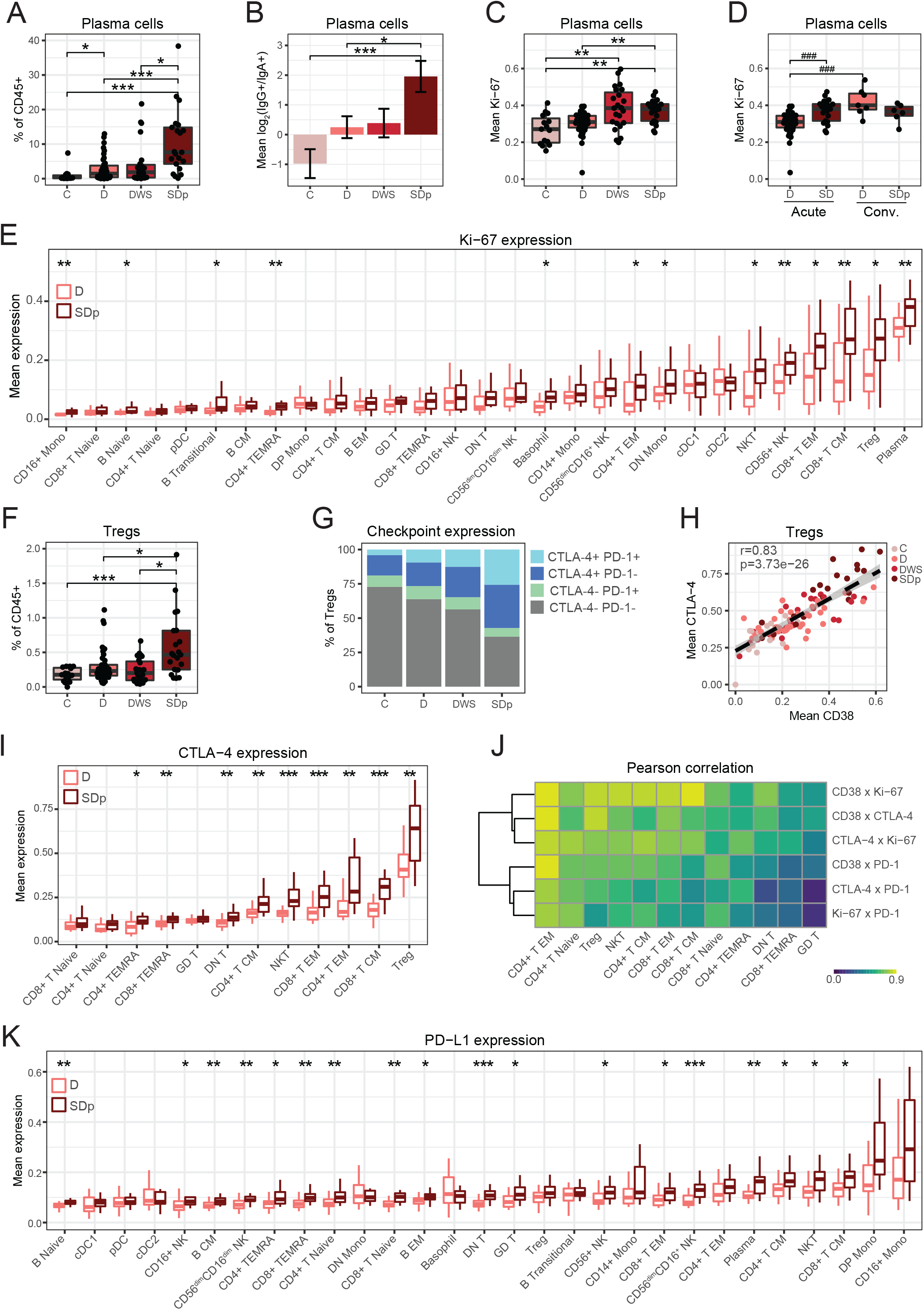
Immune activation and regulation are simultaneously enriched in acute SDp. (A, F) Box plots of plasma cell (A) and Tregs (F) fraction of CD45+ cells by clinical status. B) Mean log_2_ ratio of patient IgG^+^ plasma cell abundance to IgA^+^ plasma cell abundance by clinical status. Error bars represent standard error of mean (SEM). (C-E, I, K) Box plots of mean Ki-67 (C-E), CTLA-4(I) and PD-L1 (K) expression in acute (C-E, I, K) and convalescent samples (D) in the indicated cell populations by clinical status. (G) Proportion of CTLA-4+, PD-1+ Tregs by clinical status. Quantifications were derived from an equal subsampling of Tregs by clinical status and patient. (H) Scatter plot of mean expression of CD38 versus CTLA-4 in Tregs by patient (dots), colored by clinical status. Fitting line and CI were derived by linear regression. Pearson correlation coefficient (r) and p-value are shown. (J) Pearson correlation coefficients (color) in pairwise analysis of mean expression of checkpoint and activation molecules (rows) by T cell population (columns). All correlations are significant (q<0.05 & |effect|>0.5) except CTLA-4 x PD-1 and Ki-67 x PD-1 in GD T cells. In boxplots, center line signifies median, box signifies interquartile range (IQR) and whiskers signify IQR +/-1.5*IQR. Dots in A, C, D, F, and H represent individual patients. * q<0.05 & |effect|>0.5; ** q<0.01 & |effect|>0.5; *** q<0.005 & |effect|>0.5; # p<0.05 & |effect|>1.0; ## p<0.01 & |effect|>1.0; ### p<0.005 & |effect|>1.0 by Wilcoxon rank sum tests. Q-values represent FDR-corrected p-values. C, controls; D, dengue; DWS, dengue with warning signs; SDp, SD progressors.

Next, we asked if increased immune cell proliferation and activation was a common feature of SD progression. Thirteen of 29 cell subtypes, encompassing B, T, NK, and myeloid cell populations had higher mean Ki-67 expression in acute SDp than D samples (q<0.05) (**Figure 2E**). These trends held when quantifying Ki-67 percent positivity (**Figure S2C**). Moreover, the mean expression of the general activation marker CD38, which is involved in cell adhesion, migration, and signal transduction, among others (Hogan et al., 2019), was higher in SDp than D in 14 of 29 cell subtypes (q<0.05), confirming the greater activation of multiple cell populations via an orthogonal measurement (**Figure S2D**). The differences in Ki-67 and CD38 expression largely resolved at convalescence, although plasma cells and Tregs maintained the highest median Ki-67 expression levels in both SDp and D patients (**Figures S2E and S2F**).

Since both abundance (**Figure 1F**) and Ki-67 expression (**Figure 2E**) of Tregs were greater in SDp than D patients, we measured Treg abundance in all disease severity categories. Treg abundance in SDp was greater (median=0.47%) than DWS (median=0.20%, q=0.026), D (median=0.23%, q=0.025) and C (median=0.18%, q= 0.001) (**Figure 2F**), and these differences resolved at convalescence (**Figure S2G**). The expression of checkpoint molecules (Buchbinder and Desai, 2016) was also altered in Tregs, with higher levels of CTLA-4 in SDp than DWS (q=0.036), D (q=0.004), and C subjects (q=0.00003) and PD-1 in SDp than D (q=0.04) and C subjects (q=0.001) (**Figure 2G**). Moreover, CTLA-4 expression was highly correlated with the expression of CD38 (r=0.83, p=3.73e−26) (**Figure 2H**), which was shown to be associated with a highly suppressive Treg phenotype (Feng et al., 2017a).

To understand the breadth of checkpoint molecule upregulation in SDp, we monitored their expression in other T cell populations. CTLA-4 was more highly expressed in SDp over D in 9 of 12 T cell subtypes (q<0.05) (**Figure 2I**). PD-1 expression demonstrated similar trends, although it was significantly increased in SDp over D in only 2 of the 12 T cell populations (q<0.05) (**Figure S2H)**. As in the Tregs, we observed a significant correlation (q<0.05) between the expression of activation proteins, CD38 and Ki-67, and checkpoint proteins, CTLA-4 and PD-1, in all T cell populations (**Figure 2J**). Moreover, the checkpoint ligand PD-L1 was more highly expressed in SDp relative to D in 18 of 29 cell populations (q<0.05), spanning B, T, and NK cell populations, but not myeloid cells (**Figure 2K**). Increased CTLA-4 expression in SDp resolved in all T cell populations at convalescence **(Figure S2I)**, whereas increased PD-L1 expression persisted in some T and NK cell populations **(Figure S2J)**. These findings reveal substantial and simultaneous immune activation and regulation uniquely occurring in SDp.

### Diminished HLA-DR and increased CD64 expression on myeloid cells are hallmarks of SD progression

Next, we further probed alterations in myeloid cell abundance and function. Within the monocyte population, the fraction of CD16^+^ monocytes was lower in SDp than D (q=0.0003) and C (q=0.0008) (**Table S2**) with trends towards a reduced double negative (DN, CD14^-^CD16^-^) monocyte fraction and expanded CD14^+^ monocyte fraction which will need to be explored in larger cohorts (**Figure 3A**). Beyond differences in cell subset abundance, myeloid cells from D and SDp patients exhibited differential expression of several functional markers (**Figure 3B**). Ki-67 in CD16^+^ and DN monocytes, CD38 in these two monocyte subtypes and cDC2s, and the Fcγ receptor (FcγR) CD64 in cDC2s and DN monocytes were higher in SDp over D (q<0.05). CD141 (thrombomodulin), an anti-inflammatory factor (Li et al., 2012), was increased in CD14^+^ monocytes in SDp over D (q=0.008). A trend, albeit statistically non-significant, for greater PD-L1 expression on several myeloid cell subtypes was also detected (**Figure 3B**). In contrast, CD209 (DC-SIGN, a DENV co-receptor (Tassaneetrithep et al., 2003)) was lower on cDC2s in SDp relative to D patients (q=0.015). More prominent, however, was the reduction in HLA-DR expression in SDp relative to D patients in all myeloid cell subtypes, reaching statistical significance in DN monocytes, CD16^+^ monocytes, and cDC2s, (q<0.05) (**Figures 3B and 3C)**. Interestingly, in contrast to myeloid cells, several NK cell subtypes expressed higher levels of HLA-DR in SDp relative to D patients (q<0.05) (**Figure 3C**), suggesting increased functional activity in SDp, as previously shown in other infectious disease models (Erokhina et al., 2018; Kust et al., 2021). While the differences in proportional myeloid cell subtype abundance and HLA-DR expression largely resolved at convalescence, increased expression of CD64 in SDp over D persisted in several myeloid subsets (p<0.05) (**Figures S3A-C, E**).

**Figure 3.**
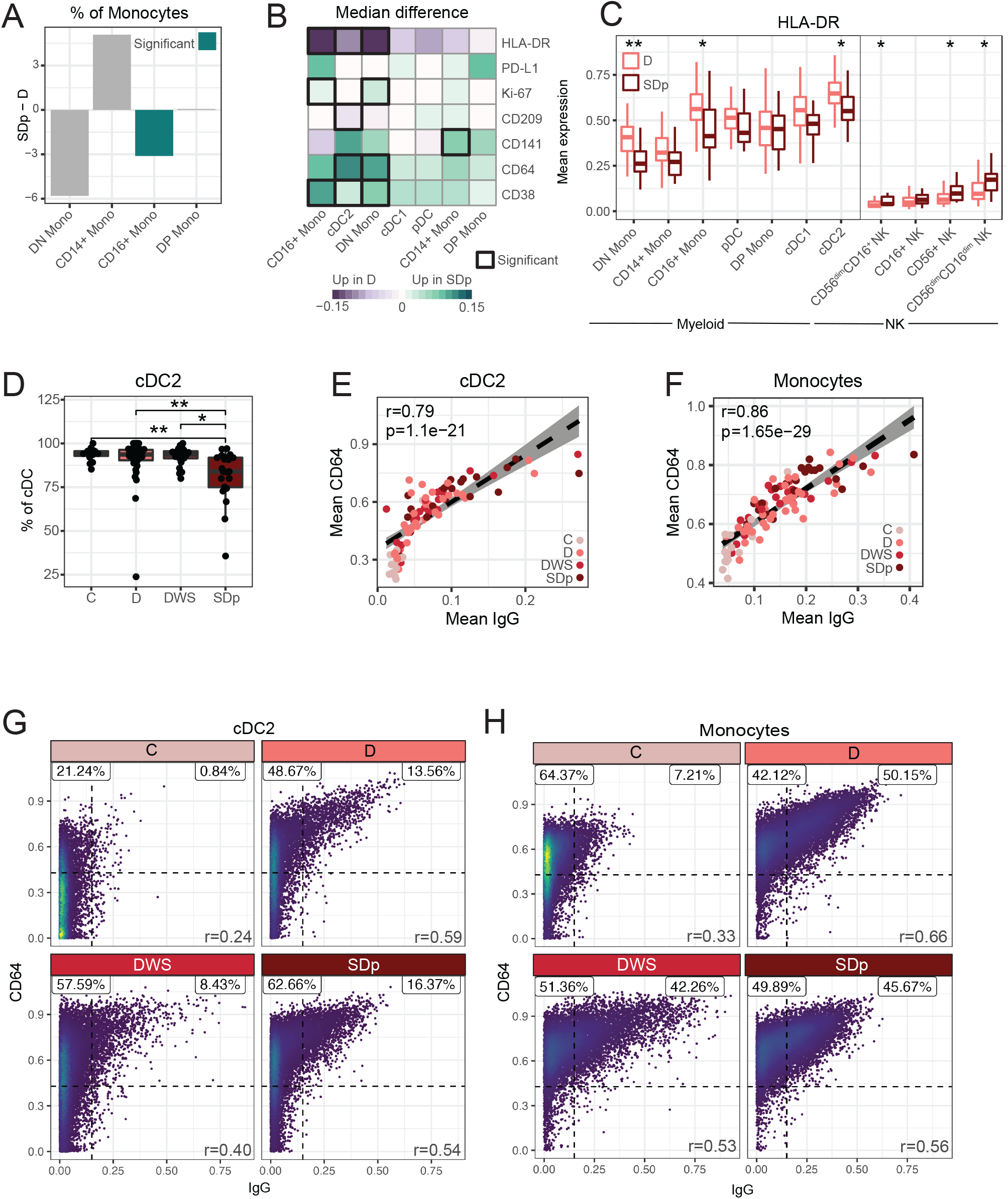
Diminished HLA-DR and increased CD64 expression on myeloid cells are hallmarks of SDp. (A) Difference in median relative abundances of monocyte subtypes (columns) between SDp and D patients out of total monocytes. (B) Difference in cohort median of patient mean expression between SDp and D across molecules (rows) in myeloid cell populations (columns). Black boxes depict significance (q<0.05 & |effect|>0.05). (C) Box plots of mean HLA-DR expression in the indicated myeloid and NK cell subtypes by clinical status. (D) Box plots of cDC2 fraction of cDCs by clinical status. Dots represent individual patients. (E-H) Scatter plots of mean (E, F) or single cell CD64 expression versus IgG detection in acute infection on cDC2s (E, G) and monocytes (F, H) by either patient (dots, colored by clinical status) (E,F) or clinical status (G, H). Fitting line and CI in E were derived by linear regression. Percent proportion of cells in each quadrant, derived from an equal subsampling of cells by clinical status and patient are shown in F and G. In boxplots, center line signifies median, box signifies interquartile range (IQR) and whiskers signify IQR +/-1.5*IQR. Teal bars (in A) and * (in C and D) indicate significance: q<0.05 & |effect|>0.5. ** q<0.01 & |effect|>0.5; *** q<0.005 & |effect|>0.5 by Wilcoxon rank sum tests. r and p-values (E, F, G) were calculated by Pearson correlation. C, controls; D, dengue; DWS, dengue with warning signs; SDp, SD progressors.

Given the changes in expression, we then asked if the proportional abundance of cDC2s out of cDCs also differed by severity. We found lower abundance in SDp than DWS (q=0.026), D (q=0.008), and C (q=.002) (**Figure 3D**), with resolution at convalescence (**Figure S3D**). Strikingly, the strongest pairwise correlation of cDC2 features appeared between CD64 expression and detected IgG (r=0.83, p=3.73e-26) (**Figure 3E**). Subsampling cDC2s from distinct disease categories revealed that at the single-cell level, CD64 and IgG were more strongly correlated in DENV-infected patients than control cells during acute infection (**Figure 3G**), whereas no such correlation was detected in convalescent samples (**Figure S3F**). Since DCs do not produce antibodies, this finding may reflect serum antibodies bound by cell surface receptors (Schatz and Ji, 2011), suggesting that CD64 expression may in part mediate the observed binding of IgG by myeloid cells. The detected level of other antibody isotypes was not correlated with CD64 expression (**Figure S3G**), in agreement with the known function of CD64 as an IgG-specific FcγR (Bruhns et al., 2009). Similar correlations between CD64 expression and IgG detection were observed on monocytes in acute samples (**Figures 3F and 3H**) and resolved at convalescence (**Figure S3H**).

Together, these findings reveal that SD progression is associated with alterations in the proportional abundance of myeloid cell populations and diminished expression of HLA-DR concurrent with increased expression of the FcγR CD64 in correlation with increased IgG antibody levels on myeloid cells, possibly mediating ADE of infection (Wang et al., 2017). Our attempts to test the latter hypothesis by measuring viral abundance in myeloid and other candidate DENV target cells (Balsitis et al., 2009; Zanini et al., 2018) were unsuccessful (**see supp material text**). Nevertheless, DENV viral load measured in patient serum via RT-qPCR was comparable between distinct disease categories in our cohort **(Figure S4A).**

### Differences in the innate immune response to DENV infection between D and SD are exaggerated in children

Since the incidence and fatality of SD are greater in children than adults (Hammond et al., 2005), we compared cell population abundances and functional marker expression by distribution tests between D and SDp adults and children (under 17 years) (**Table S4**). The abundance of cytotoxic CD56^dim^CD16^+^ NK cells was significantly lower in SDp relative to D children (p=0.002), but not adults (p=0.42) (**Figures 4A and 4B**). Moreover, the expression of the FcγR, CD16, was lower on CD56^dim^CD16^+^ NK cells in SDp children (p=0.002), but not adults (p=0.16), suggesting receptor shedding indicative of cell activation (Srpan et al., 2018) (**Figure 4C**). The drop in CD56^dim^CD16^+^ NK cells in SDp children was accompanied by a proportional expansion of CD56^+^ NK cells out of total NK cells (p=0.014) **(Table S4,** **Figure 4D**), suggesting that NK cells may be skewed towards a less cytotoxic, more inflammatory phenotype (Erokhina et al., 2018). The fraction of CD56^dim^CD16^+^ NK cells out of total NK cells was almost sufficient to stratify the cohort by disease severity in children, but not adults (**Figures 4D** **and S5A**). HLA-DR was more highly expressed on CD56^dim^CD16^+^ NK cells in SDp children but not adults, further supporting increased functional activity of these cells (Erokhina et al., 2018; Kust et al., 2021) (**Figure 4E**).

**Figure 4.**
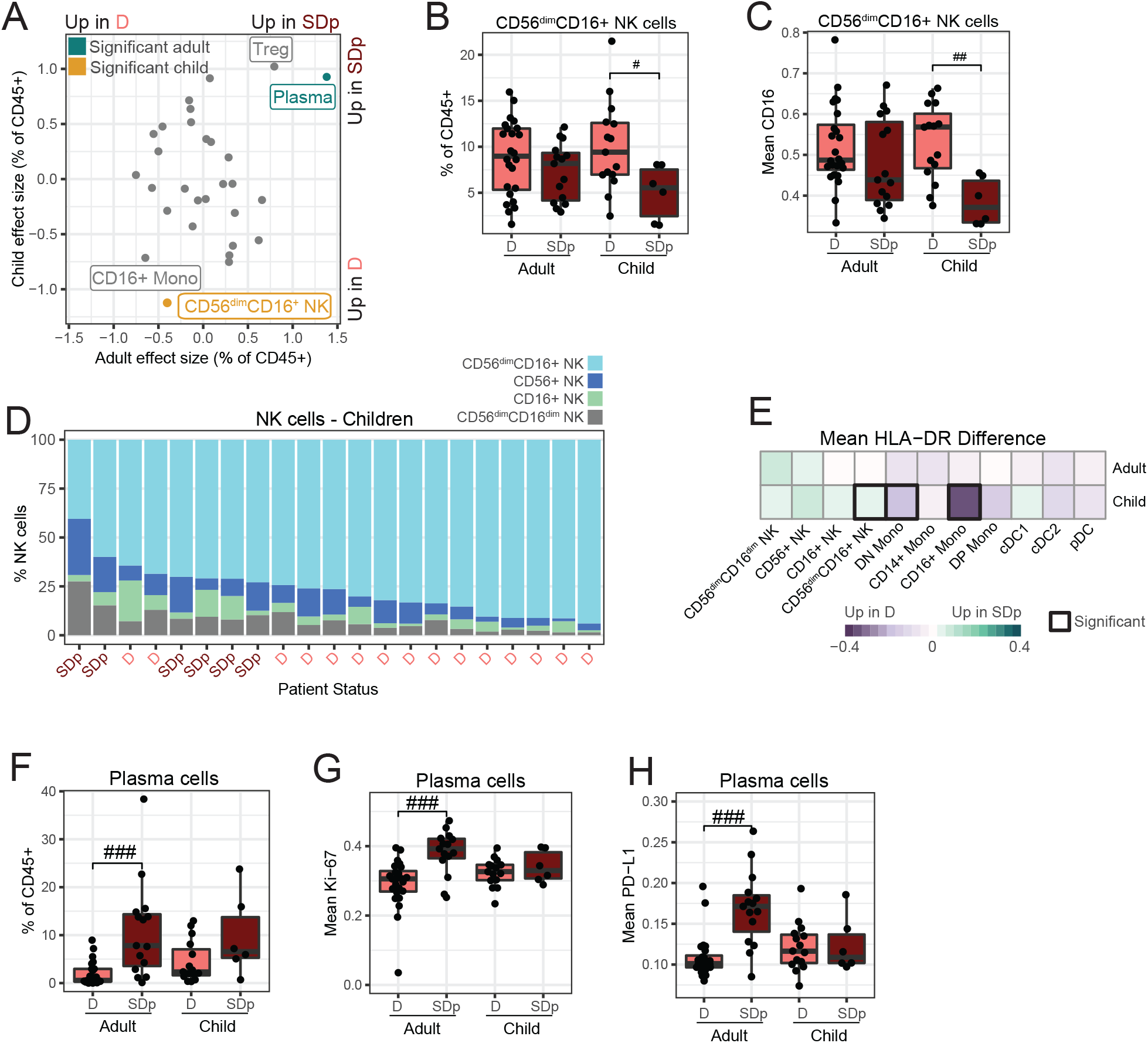
Differences in the innate immune response to DENV infection in D and SDp are exaggerated in children. (A) Effect size in comparison of cell population abundances (dots) in SDp and D in adults (x-axis) and children (y-axis). Teal and gold indicate significance in adults and children, respectively. (p<0.05 & |effect|>1.0). (B, F) Box plots of CD56^dim^CD16^+^ NK cell (B) and plasma cell (F) fractions of CD45+ cells by clinical status and age. Dots represent individual patients. (C, G, H) Box plots of mean CD16 (C), Ki-67 (G) and PD-L1 (H) expression in the indicated cells by clinical status and age. Dots represent patients. (D) Proportion of NK cell subtypes in children ordered by CD56^dim^CD16^+^ NK cell abundance. Columns represent individual patients, labeled by clinical status. (E) Difference in cohort mean of patient mean HLA-DR expression between SDp and D across cell populations (columns) by age (rows). Black boxes indicate significance (p<0.05 & |effect|>1.0). In boxplots, center line signifies median, box signifies interquartile range (IQR) and whiskers signify IQR +/-1.5*IQR. # p<0.05 & |effect|>1.0; ## p<0.01 & |effect|>1.0; ### p<0.005 & |effect|>1.0 by Wilcoxon rank sum tests. D, dengue; SDp, SD progressors.

The abundance of CD16^+^ monocytes was lower in SDp than D patients in both adults and children (**Figures 4A** **and S5B).** Nevertheless, HLA-DR was lower on CD16^+^ and DN monocytes in children only (**Figure 4E**), despite the smaller sample size, suggesting a greater impairment in antigen presentation in SDp children than SDp adults.

Plasma cell abundance appeared higher in SDp than D in both adults and children, but the difference was statistically significant in adults only (**Figures 4A and 4F**). While this difference could in theory be explained by the smaller number of pediatric samples analyzed, the effect size was greater in adults (1.38) than in children (0.93). Moreover, the expression levels of Ki-67 and PD-L1 in plasma cells were higher in SDp adults relative to D adults, but comparable in SDp and D children (**Figures 4A and 4G-H**), suggesting that the observed differences in plasma cell responses between adults and children may be biologically relevant.

While CD8^+^ TEMRAs abundance was reduced in SDp in the combined cohort (**Figure 1F**), this difference did not reach statistical significance in the distinct age groups (**Figures 4A** **and S5C**). Nevertheless, HLA-DR and CD38 were more highly expressed in CD8^+^ TEMRAs in SDp children (p<0.05), but not adults (**Figures S5D and S5E**), signifying greater cell activation. In contrast, neither abundance of (**Figures 4A, S5B and S5F)** nor expression markers (**Table S4**) on CD16^+^ monocytes and Tregs were significantly altered between SDp adults and children.

These findings reveal both overlapping and distinct immune responses between SDp adults and children and point to increased plasma cell abundance and proliferation in SDp adults, yet more inadequate expansion of activated cytotoxic NK cells, and more prominent CD8^+^ TEMRAs activation and reduced antigen presentation on myeloid cells in SDp children.

### Immune cell composition stratifies patients by SD categories

The 2009 WHO definition of SD includes the presence of severe bleeding or plasma leakage, somewhat similar to the prior definitions of DHF and DSS, respectively, but also organ impairment (OI) involving the liver, brain and heart (WHO, 2012). To compare the early host response to DENV between these SD categories, we measured cell type abundance and protein expression in patients who once progressed to SD manifested with hemorrhages and/or vascular leak (DHF/DSS, n=15) and those with organ (liver) impairment (OI, n=4) **(Figure S6, Table S5).** Children and adults were similarly represented in these two groups (**Figure S6**). Two patients who presented with both DHF and OI were not included in this analysis. The abundance of immune cell populations was dramatically altered between DHF/DSS and OI patients, with higher abundances of cDC1s, cDC2s, and Tregs in OI, yet higher abundance of plasma cells in DHF/DSS (p<0.05) (**Figure 5A**).

The abundances of cDC1s and cDC2s in OI resembled and were not statistically different from C (p=0.36 and p=0.18). cDC1 and cDC2 abundances were, however, lower in DHF/DSS compared to C (p=0.0001 and p=0 .000001) (**Figure 5B**). In fact, ordering patients by total cDC abundance nearly perfectly stratified the two SD categories (**Figure 5C**). Additionally, cDC1s from OI samples expressed a higher Ki-67 level (p=0.01), but lower CD64 level (p=0.02) (**Figure 5D**), suggesting that while expanded and more proliferative relative to DHF/DSS, they may have decreased antibody binding capacity.

**Figure 5.**
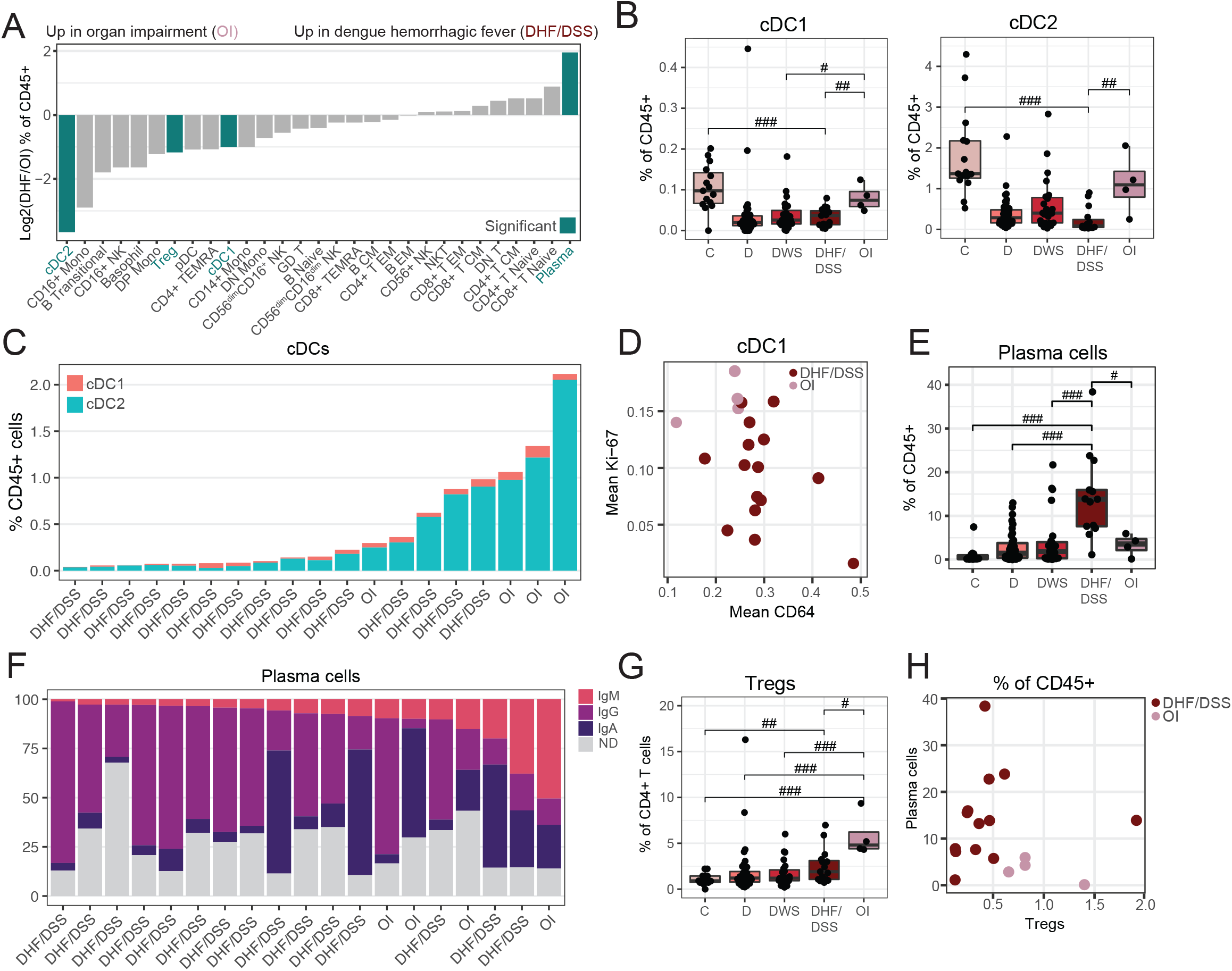
Immune cell composition stratifies patients by SD categories. (A) Log_2_ ratio of median abundances of cell populations (columns) between patients with DHF/DSS and OI. Teal bars indicate significance (p<0.05 & |effect|>1.0). (B, E, G) Box plots of cDC1 (B, left), cDC2 (B, right), plasma cell (E), and Treg (G) fractions of CD45+ cells by clinical status. Dots represent individual patients. (C) Proportion of cDC populations ordered by total abundance. Columns represent individual patients, labeled by SD syndrome. (D) Mean CD64 and Ki-67 expression in cDC1s by patient (dots), colored by SD syndrome. (F) Proportion of isotype usage in plasma cells ordered by IgM usage. Columns represent individual patients labeled by SD syndrome. ND, isotype not determined due to low expression. (H) Scatter plot of plasma cell and Treg abundance by patient (dots), colored by SD syndrome. In boxplots, center line signifies median, box signifies interquartile range (IQR) and whiskers signify IQR +/-1.5*IQR. # p<0.05 & |effect|>1.0; ## p<0.01 & |effect|>1.0; ### p<0.005 & |effect|>1.0 by Wilcoxon rank sum tests. OI, organ impairment; DHF/DSS, Dengue hemorrhagic fever/dengue shock syndrome.

Beyond myeloid cells, plasma cell abundance was greater in DHF/DSS than OI (and C, D, DWS) patients (p<0.05) (**Figure 5E**). The immunoglobulin isotype usage in plasma cells was also different, with predominately IgG enrichment in DHF/DSS patients vs. an increase in IgM enrichment in OI (p=0.04), revealing a less class-switched, more naïve humoral response in OI (**Figure 5F**). In contrast, Tregs were more abundant in OI than DHF/DSS, DWS, D, and C samples (p<0.05) (**Figure 5G**), yet there was no significant difference in CTLA-4 and PD-1 expression between the two SD categories (**Table S5**). OI is thus skewed towards Tregs, whereas DHF/DSS is skewed towards plasma cells. Indeed, plotting these two features on a biaxial plot nearly perfectly stratified the two SD categories (**Figure 5H**). Although numbers are small, these differences in immune cell composition and activation support that DHF/DSS and OI may represent distinct syndromes associated with differential immunopathogenesis.

### The temporal switch of innate and adaptive immune activation and concurrent immune regulation is dysregulated in acute SDp

To capture the temporal dynamics in the host response during acute infection, we monitored the kinetics of immune activation and regulation in SDp and D on days 3 to 8 post fever onset (**Figures S7A and S7B**). For each individual day, we used samples from a 3-day rolling window (**Table S1.4 and Table S6**) to account for error in the timing of patients’ self-reported symptom onset. We performed Wilcoxon rank sum tests and calculated effect size between SDp and D for each rolling day window for all features (**Table S6**).

Given the significant increase in immune activation seen in SDp over D (Fig. 2E, S2D), we asked if this difference varied throughout the course of infection. We assessed the difference in mean expression of the activation marker, CD38, by time and cell population and found significant differences in T and myeloid cell populations (p<0.05) (**Figure 6A**). Three clusters emerged when the data were hierarchically clustered by cell population: populations with higher CD38 expression in SDp early in acute infection, populations with higher CD38 in SDp late in acute infection, and populations with similar levels of CD38 at all timepoints. The first cluster consisted primarily of memory T cell populations, whereas the second cluster was comprised primarily of myeloid populations.

**Figure 6.**
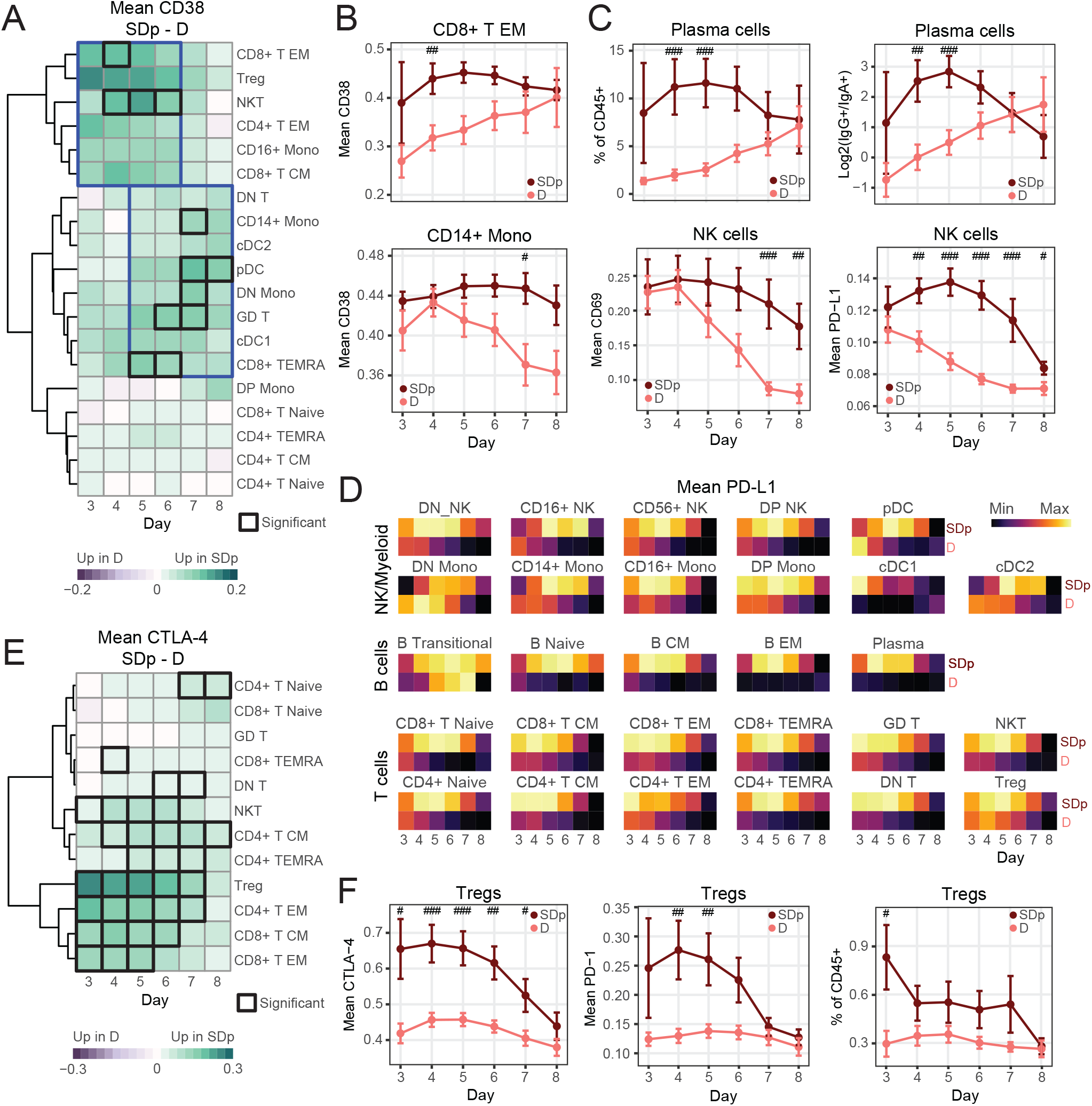
The temporal switch of innate and adaptive immune activation and concurrent immune regulation is dysregulated in SDp. (A, E) Difference in cohort mean of patient mean CD38 (A) and CTLA-4 (E) expression between SDp and D across cell populations (rows) by day (columns). Black boxes depict significance (p<0.05 & |effect|>1.0). Blue boxes group timepoints and populations with similar trends. (B) Cohort mean of patient mean CD38 expression in CD8^+^ T EM (top) and CD14^+^ monocytes (bottom) by clinical status and day. (C) Cohort mean of plasma cell abundance (top center), log_2_ ratio of IgG^+^ to IgA^+^ plasma cell abundance (top right), and mean CD69 (bottom left) and PD-L1 (bottom right) expression in NK cells by clinical status (color) and day (columns). (D) Cohort mean of patient mean PD-L1 expression (color) by day (columns) and clinical status (rows), separated by cell population (tiles). Each tile is individually scaled. (F) Cohort mean of patient mean CTLA-4 (left) and PD-1 (center) expression in Tregs and abundance of Tregs (right) by clinical status and day. # p<0.05 & |effect|>1.0; ## p<0.01 & |effect|>1.0; ### p<0.005 & |effect|>1.0 by Wilcoxon rank sum tests. Error bars represent SEM.

To better understand these dynamics, we plotted the mean expression of CD38 in D and SDp over time in a representative cell population from each of the first two clusters: CD8^+^ T EM (higher early in SDp) and CD14^+^ monocytes (higher late in SDp) (**Figure 6B**). In CD8^+^ T EM cells, while CD38 was continuously more highly expressed starting on day 3 in SDp, its expression in D patients was low early on but gradually increased to a similar level as SDp by day 8, with the difference peaking on day 4 post fever onset. This early CD8^+^ T EM cell activation could be explained in part by the higher percentage of secondary infections among SDp (**Figure S1A**). Contrastingly, in CD14^+^ monocytes, CD38 mean expression was initially high in both D and SDp, yet while it steadily declined over the course of acute infection in D, in SDp, it remained high throughout the acute infection, with the difference peaking on day 7. CD64 expression on myeloid cells followed similar trends to CD38, with significantly higher expression in SDp in cDC2s and DN monocytes at late acute timepoints (p<0.05) (**Figure S7C**). These shifts in the kinetics of the adaptive and innate immune responses were not limited to T and myeloid cells, but also appeared in plasma cell abundance and isotype usage, as well as the expression of the stimulatory receptor, CD69 (Borrego et al., 1999), on NK cells (**Figure 6C**).

Next, we monitored the kinetics of immune regulation during the course of acute DENV infection. PD-L1 expression on NK cells was comparable in D and SDp patients on day 3, yet in D patients, it steadily declined through day 8, whereas in SDp, its expression peaked on day 5, an intermediate timepoint between the first and second clusters, before diminishing (**Figure 6C**). Strikingly, this pattern was conserved in nearly every immune cell population (**Figure 6D**), though the magnitude of PD-L1 expression varied (**Figure S7D**). This temporal pattern of PD-L1 expression was perturbed only in transitional B cells, which peaked at day 7 in D, rather than early in infection. This antigen-inexperienced population has been shown to be enriched for immunoregulatory cytokine production (Glass et al., 2022), which may coincide with their unique checkpoint expression profile. Interestingly, the CTLA-4 expression pattern was more similar to that of CD38 (**Figures 6A and 6E**), with higher expression in Tregs, CD4^+^ T EM, CD8^+^ T EM, and CD8^+^ T CM in SDp primarily at earlier timepoints (p<0.05) (**Figure 6E**). This temporal coordination between CTLA-4 and CD38 in T cells was consistent with a high degree of correlation measured between these two markers (**Figures 2H and 2J**). CTLA-4 and PD-1 expression levels in Tregs were higher in SDp at early timepoints (p<0.05) before gradually decreasing, whereas in D patients, both proteins were expressed at relatively lower levels throughout infection (**Figure 6F**). Likewise, the overall Treg abundance peaked early in SDp (p<0.05), while it remained stably lower in D.

HLA-DR showed no obvious temporal patterns, yet it was significantly lower in multiple myeloid cell populations in SDp relative to D patients throughout the disease course (p<0.05) (**Figure S7E**). Moreover, the proportional abundance of CD16^+^ and DN monocytes, on which HLA-DR expression was lower in SDp, was low in SDp throughout the infection, whereas in D patients, it started high and gradually decreased (**Figure S7F**). The abundance of CD56^dim^CD16^+^ NK cells was also lower in SDp than D during acute infection (**Figure S7G**).

Based on these findings, we propose a model in which uncomplicated D is characterized by early activation of the innate immune response predominated by expansion of CD16^+^ monocytes, which along with other myeloid cells, present antigen adequately via HLA-DR expression, as well as increased DENV target cell killing via cytotoxic NK cells. This response then wanes as the adaptive immune system becomes increasingly stimulated, with minimal activation of regulatory signals. Contrastingly, in SDp, there is rapid expansion of IgG-producing plasma cells (more pronounced in adults) and memory T cell activation (more pronounced in children). Moreover, a more activated Treg population emerges concurrently with inadequate expansion of CD16^+^ monocytes and reduced antigen presentation capacity in myeloid cells (particularly in children) as well as skewing of NK cell responses from cytotoxic to proinflammatory (particularly in children). Together, this uncoordinated immune response is ineffective in controlling viral spread and at the same time increases proinflammatory signals causing tissue injury and leading to worse prognosis (**Figure 7**).

**Figure 7.**
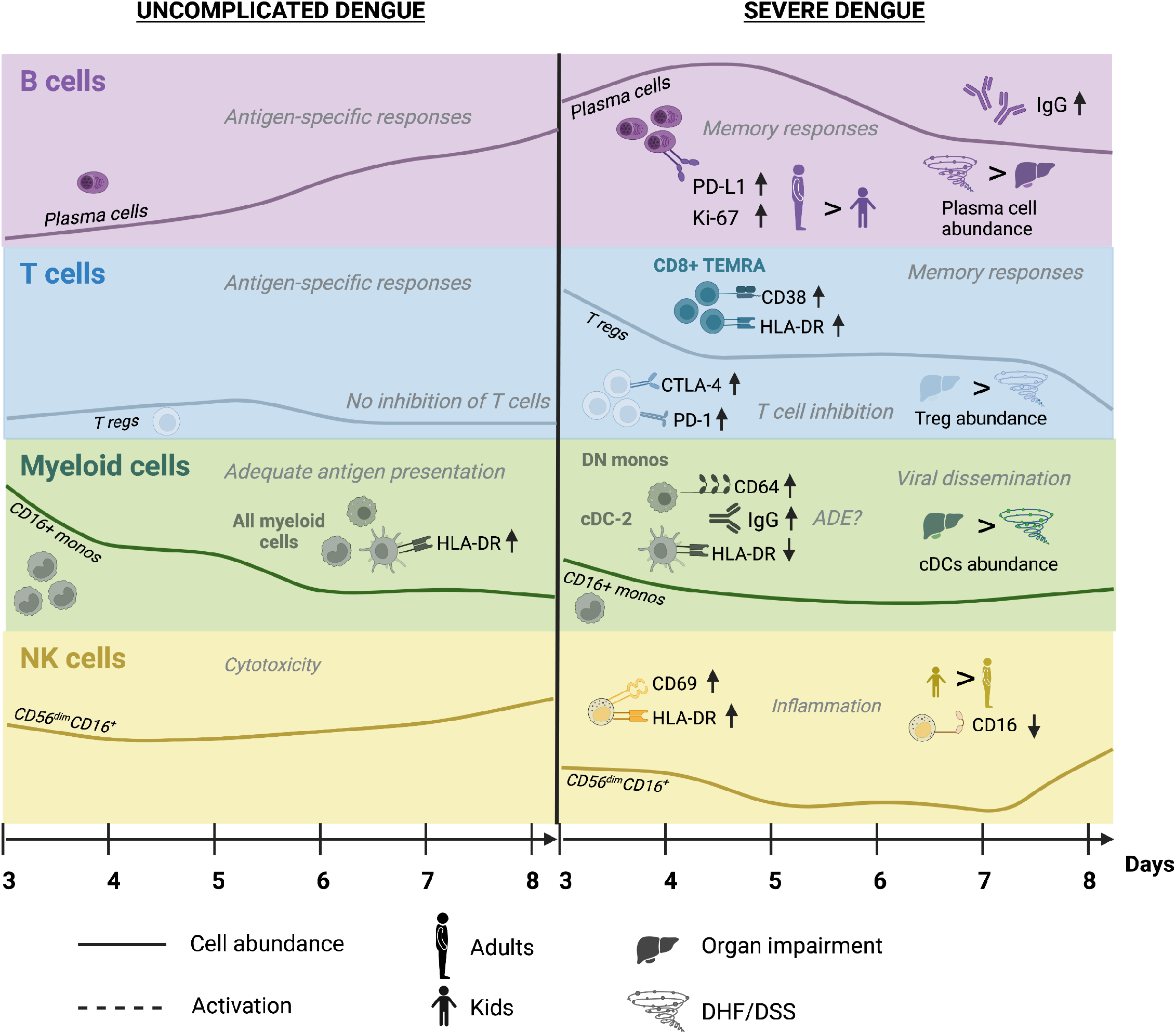
Proposed model for differential immune responses between uncomplicated and SD patients. Schematic of the kinetics and magnitude of cell subtype abundance and protein marker expression during acute D and SD infections. Most prominent differences in innate and adaptive immune responses are shown. Prominent differences in immune responses between adults and children and SDp with DHF/DSS and OI are also depicted.

## Discussion

The pathogenesis of SD remains poorly understood. Here, we comprehensively characterized the human immune response to DENV infection at single-cell resolution. Leveraging the clinical diversity of the Colombia dengue cohort to capture real-world heterogeneity revealed unique and overlapping responses between various disease categories and age groups during the course of natural infection.

Previous studies using similar approaches to monitor natural DENV infection have uncovered increased type I IFN responses (Zhao et al., 2020), upregulation of activation markers including CD38 and CD169 on myeloid cells (Fenutria et al., 2021), and temporal shifts in B and T cell phenotypes (Rouers et al., 2021). However, these studies either did not include SD patients, combined SD with D patients in comparison with controls, or studied focused elements of the immune response. Several features of our cohort and workflow have provided a unique opportunity to probe SD pathogenesis. First, our study included 22 SD patients, 19 of whom were enrolled at an early disease stage, enabling monitoring of the host response preceding the progression to SD. Second, enrolling adults and children enabled identification of age-dependent and independent determinants in the host response to DENV. Third, comparing SDp manifesting with OI to those with DHF/DSS has shone light on this previously unstudied aspect, defining these conditions as likely immunologically distinct syndromes. Beyond clinical heterogeneity, we captured immune heterogeneity using an antibody panel that enabled clustering of a diverse array of cell subtypes. Lastly, applying this analysis to longitudinal patient samples collected during the disease course and at convalescence facilitated a comprehensive profiling of both the magnitude and kinetics of the immune responses to DENV infection.

Our findings revealed greater and more rapid activation of a memory response in SDp than D patients. A prominent expansion of proliferating IgG-secreting plasma cells, as previously described (Appanna et al., 2016; Kwissa et al., 2014), was measured early in the course of SDp, whereas the magnitude of this response was lower and its kinetics slower in D patients. Similarly, SDp, particularly children, demonstrated increased CD38 and HLA-DR expression on CD8^+^ T EM and/or TEMRA cells, suggesting more robust activation of mature T cell responses. These findings support the original antigenic sin theory in which pre-existing memory B and T cells generated from a prior heterotypic DENV infection skew the early immune response upon secondary infection towards a less specific yet more pro-inflammatory phenotype leading to disease progression (Rothman, 2011).

Monocyte subsets from SDp expressed higher levels of CD38, whose binding to CD31 on platelets and endothelial cells has been shown to facilitate adhesion and migration across the endothelial barrier (Feng et al., 2017b), suggesting a possible role in promoting inflammation and/or vascular leakage. Concurrent with increased inflammatory signals, our findings provide evidence for impaired antigen presentation by myeloid cells in SDp. While CD16^+^ monocyte abundance in D was maintained at similar levels to controls, in SDp, despite increased Ki-67 expression, their abundance was lower than in D. This inadequate expansion of cells with antiviral functions suggests potential impairments in scavenging infected cells, neutrophil recruitment and patrolling leading to reduced viral clearance and antigen presentation in SDp (Buscher et al., 2017; Marsh et al., 2017). The lower expression of HLA-DR on CD16^+^ (and DN monocytes and cDC2s) in SDp provides additional evidence for impaired antigen presentation by these myeloid cell subtypes.

Since CD16 (type III FcγR) facilitates phagocytosis and is implicated in ADE of DENV infection (Wang et al., 2017), the drop in CD16^+^ monocyte abundance, may also reflect infection-induced cell death. The reduced proportional abundance of cDC2s (out of cDCs) in SDp suggests possible target cell killing or apoptosis and DENV enhancement in cDC2s. Indeed, CD64 (high affinity type I FcγR) was more highly expressed in SDp particularly on cDC2s, and its expression level was highly correlated with detected IgG on these cells, suggesting a role in ADE. This finding appeared specific to CD64, as the expression of CD32, another activating FcγR implicated in ADE (Bournazos et al., 2020; Chawla et al., 2013; Rodrigo et al., 2009), was comparable across severity categories. It is also tempting to speculate that the lower expression of the DENV co-receptor CD209 on cDC2s resulted from virus-receptor complex internalization.

These findings agree with prior knowledge that cDC2s, but not cDC1s are permissive to infection with other enveloped, endocytic viruses (Silvin et al., 2017). Our findings propose candidate specific cell types and receptors which drive ADE of infection. Nevertheless, since our attempts to detect viral proteins via mass cytometry were unsuccessful, we were unable to identify DENV target cells and explore functional differences between infected and bystander cells. It is likely that high viral titers combined with viral components shed into plasma made the detection of genuinely infected dengue cells indistinguishable from the background milieu (**Figure S4D**).

Our findings revealed a dramatic expansion of Tregs in SDp only, which may be driven in part by the rapid activation of myeloid and plasma cells. The higher expression of CD141, an anti-inflammatory factor typically expressed on cDC1s, on CD14^+^ monocytes in SDp may enhance this Treg response. Indeed, CD141 expression on circulating monocytes, was shown to induce Treg responses in other disease models (Chu et al., 2012; van Leeuwen-Kerkhoff et al., 2020). HLA-DR^low^ CD14^+^ monocytes, a regulatory myeloid cell subtype which trended towards expansion in SDp may also promote Treg expansion, as shown in other viral infections (Ren et al., 2016; Wang et al., 2016).

Tregs were previously reported to expand in D patients relative to healthy controls (Lühn et al., 2007) (Jayaratne et al., 2018), yet their role in disease progression remained unclear. Whereas a previous study demonstrated a protective effect via inhibition of vasoactive cytokines (Lühn et al., 2007), another showed no correlation between disease severity and a less suppressive Treg phenotype (Jayaratne et al., 2018). Our findings that SDp Tregs display significantly higher levels of the effector molecules CTLA-4, PD-1 and CD38 than D Tregs, suggest a more suppressive phenotype associated with SD (Feng et al., 2017a). We observed similar findings in other T cell populations, indicating a globally inhibited T cell response which may be associated with SD progression. The lower expression of HLA-DR on myeloid subtypes in SDp provides additional evidence for the association of SD with suppressed T cell stimulation via impaired antigen presentation.

On one hand, a suppressive Treg response could be detrimental early in the disease course by inhibiting protective antiviral responses via apoptosis of antigen-specific T cells concurrent with inhibition of Treg cell death (Pandiyan et al., 2007; Pierson et al., 2013), thereby extending uncontrolled viral replication and enhancing pathology. Indeed, increased expression of CTLA-4 and PD-L1 on T cells has been associated with Ebola virus disease fatality (Ruibal et al., 2016), and polymorphisms in the CTLA-4 gene are associated with SD progression (R.-F. Chen et al., 2009). On the other hand, suppression of proinflammatory signals may protect from cytokine storm and/or tissue injury. Interestingly, Treg activation peaked early in SDp and started to gradually decline prior to the onset of SD, suggesting that it may provide some protection. Moreover, the greater abundance of Tregs in SDp presenting with OI than DHF/DSS also suggests that Tregs may protect from cytokine storm.

While most immune alterations in SDp largely resolved at convalescence, among the few that persisted were higher expression of PD-1 on Tregs and EM CD4^+^ T cells and PD-L1 on NK cells. Sustained checkpoint protein expression may enhance immune regulation upon subsequent infection (Simon-Lorière et al., 2017), and/or represent a pre-existing immune setpoint altering dengue severity.

SD diagnosis based on the more recent 2009 WHO criteria includes evidence for OI, in addition to plasma leakage and bleeding (DHF/DSS) (WHO, 2012). Nevertheless, in part due to the limited number of SD cases, no prior studies have compared these conditions, and it remained unknown whether they represented a continuous disease spectrum or distinct syndromes with differing pathogenic mechanisms. We provide evidence that while largely overlapping, some differential responses between SDp presenting with DHF/DSS or OI exist. First, the abundance of cDC2s, which expressed higher CD64 in correlation with detected IgG and expressed lower CD209, was greater in OI than DHF/DSS. It is tempting to speculate that cDC2s serve as Trojan horses for DENV, facilitating dissemination to organs and tissue injury. The expansion of Tregs in SDp was also driven by OI SDp, further supporting the hypothesis that reduced DENV clearance facilitating dissemination plays a role in this SD phenotype. Beyond viral dissemination, it was shown that CD8 T cell responses are implicated in liver injury in the setting of DENV infection (Sung et al., 2012). Intriguingly, cDC1s, which stimulate such CD8^+^ T cell responses, were expanded in OI but not DHF/DSS SDp. In contrast, plasma cell abundance and IgG usage were more prominent in DHF/DSS than OI, suggesting that ADE and/or another mechanism, such as complement activation in response to immune complex formation, may promote secretion of vasoactive products leading to bleeding and/or shock (Ricklin and Lambris, 2013; Xu et al., 2012). Production of vasoactive cytokines may be further augmented by the reduced Treg expansion in DHF/DSS patients, in contrast to OI SDp (Lühn et al., 2007). It will be important to monitor these responses in a larger number of SDp and in infections involving other organs beyond liver.

Evidence for NK cell activation in SDp was detected in the combined cohort, with SDp showing higher expression of Ki-67 and CD38, and in agreement with a former study, CD69, an activating receptor that contributes to sustained NK-cell cytotoxicity, proliferation and TNF-α and adhesion molecule production (Borrego et al., 1999; Green et al., 1999). Nevertheless, comparing children and adult NK cell responses highlighted additional findings. On one hand, CD56^dim^CD16^+^ NK cells, which have known cytotoxic functions, showed evidence of increased activation in SDp children relative to SDp adults. First, the expression level of CD16 in SDp children was lower than SDp adults. Since CD16 has been shown to be shed by metalloproteinases upon NK cell activation, thereby increasing NK cell survival and cytotoxicity via serial engagement of target cells (Srpan et al., 2018), this finding suggests that these cells are activated. Thus, the predicted impact of this finding on clearance of DENV-infected cells is unclear. Second, CD56^dim^CD16^+^ NK cells in SDp children but not adults expressed higher HLA-DR. Interestingly, HLA-DR expression on NK cells has been implicated in other disease models not only in proliferation, degranulation, and IFN-γ production, but also in antigen presentation and activation of T cell responses (Erokhina et al., 2018; Kust et al., 2021). It is tempting to speculate that NK cells are activated, at least in part, to compensate for the lack of adequate T cell responses, thereby promoting viral clearance and protection.

On the other hand, it appears that in SDp, and particularly SDp children, such compensation may not be sufficient/effective, since the abundance of the CD56^dim^CD16^+^ NK cell population was reduced relative to D patients. This finding uncovers a potential reduction not only in NK cell-mediated killing of virally-infected cells, but also NK cell homing to tissues (Béziat et al., 2011; Castriconi et al., 2018) (Zimmer et al., 2019). The concurrent expansion of cytokine producing CD56^+^ NK cells suggests that enhanced proinflammatory responses in SDp children relative to adults may further increase vascular permeability, more commonly reported in children (Guzmán et al., 2002).

Beyond altered NK cell responses, SDp children demonstrated greater reduction in HLA-DR expression on myeloid cell populations than SDp adults and a trend towards reduction of CD16^+^ monocytes, suggesting a greater impairment in antigen presentation and possibly other monocyte functions. Moreover, CD8^+^ TEMRA in SDp children, but not SDp adults, expressed higher levels of HLA-DR. While there was no difference in checkpoint molecule expression on this cell type between disease severity categories, the increase in CD38 expression detected in children, may suggest a more activated CD8^+^ T cell response. In contrast, SDp adults showed a more prominent expansion of proliferating plasma cells expressing high levels of PD-L1 than SDp children. Together, these findings provide evidence that suppressed innate immune responses involved in cell killing and antigen presentation impairing viral clearance, and increased activation of CD8^+^ TEMRAs may predominate in SDp children, whereas plasma cell expansion predominates in SDp adults. Nevertheless, since the fraction of patients with secondary infection was greater in adults than children (76% vs. 53%), we cannot exclude a possibility that differences in dengue exposure beyond age account for some of the observed differences.

A limitation of our study is that our cohort included more SDp females than males (16 vs. 6). Since female gender and pregnancy are reported risk factors for SD progression (Machado et al., 2013; Sangkaew et al., 2021), monitoring these responses in a gender-balanced cohort is important. We were also motivated to compare the immune response in SDp who had no prior exposure to DENV (primary infection), where ADE is unlikely to play a role, with secondarily infected SDp. Nevertheless, the number of definitive primary SDp was low (n=2). Moreover, genetic background plays a role in SD pathogenesis, as evidenced by the lower incidence of SD progression in Africa (Gainor et al., 2022). Therefore, probing the generalizability of our findings in other cohorts and identifying cell-specific generalizable signatures, as those we have recently identified in bulk samples (Liu et al., 2022; Robinson et al., 2019), is also important. Lastly, future studies are needed to capture the host response following the mosquito bite and during the first 2-3 days of symptoms, yet this can only be achieved through inoculation studies which pose ethical challenges.

There is an urgent need for a prognostic assay to predict SD early in the disease course to improve patient triage and help select patients for therapeutic studies. We and others have recently identified candidate clinically usable biomarkers (Bournazos et al., 2021; Liu et al., 2022; Robinson et al., 2019; Tissera et al., 2017; Zanini et al., 2018). Yet, additional biomarkers are needed until a prognostic assay is available and validated. Changes in cell subtype abundance and/or marker expression that precede the development of SD, as those identified in this study, could serve as candidate predictive biomarkers. Moreover, these determinants may serve as potential drug targets to prevent progression to SD. While speculative, CD38 inhibitors (Feng et al., 2017a; Mauriello et al., 2020) and CTLA-4 and PD-1 inhibitors (Wykes and Lewin, 2018) are a few examples for existing strategies that based on our data may modulate immune responses to favor viral clearance. Indeed, blockade of PD-1/PD-L1 interaction with a monoclonal antibody in PBMCs isolated from patients infected with another Flaviviridae, hepatitis C virus, restored exhausted CD8 T cell responses leading to increased T cell proliferation and cytokine production (Golden-Mason et al., 2007; Urbani et al., 2006). Further validation and functional characterization of these candidate biomarkers and therapeutic targets is required.

In summary, this study provides insight into the pathogenesis of SD in natural infection in humans at a high resolution and proposes cellular and molecular determinants as candidate predictive biomarkers and/or targets for countermeasures to prevent SD.

## Methods

### Study population

PBMC samples were collected from individuals presenting to the Fundación Valle del Lili in Cali, Colombia or Centro de Atención y Diagnóstico de Enfermedades Infecciosas (CDI) in Bucamaranga between 2016 and 2019 with symptoms compatible with dengue as previously described (Liu et al., 2022). All work with human subjects was approved by the Stanford University Administrative Panel on Human Subjects in Medical Research (Protocols #35460 and #50513) and the ethics committees in biomedical research of the Fundación Valle del Lili (Cali/Colombia) and the Centro de Atención y Diagnóstico de Enfermedades Infecciosas (CDI) (Bucaramanga/Colombia). All subjects, their parents, or legal guardians provided written informed consent, and subjects between 6 to 17 years of age and older provided assent.

### PBMC isolation

PBMCs were isolated using SepMate tubes (Stemcell Technologies) according to the manufacturer’s instructions. Whole blood was diluted 1:1 with phosphate-buffered saline (PBS) and added to a SepMate tube, which contained 15 ml of Ficoll. Tubes were then centrifuged for 10 minutes at 1,200g, after which the PBMC layer was poured off into a fresh tube and washed with PBS. Tubes were then centrifuged at 250 x g for 10 minutes and resuspended in freezing media. Cryovials containing PBMCs were then placed in a CoolCell at -80°C for 24 hours prior to being transferred to liquid nitrogen for storage. Samples were also shipped in liquid nitrogen.

### Confirmation of dengue diagnosis

The confirmation of dengue diagnosis and screening for other arboviruses including Zika virus and chikungunya virus was performed as previously described (Robinson et al., 2019). Briefly, serum samples were screened with a qualitative, single-reaction, multiplex real-time reverse transcriptase polymerase chain reaction (rRT-PCR) that detects Zika, chikungunya, and DENV RNA (Waggoner et al., 2016). The DENV serotype and viral load were detected and quantitated using a separate DENV multiplex rRT-PCR (Waggoner et al., 2013).

Serologic and avidity testing were performed via a multiplexed antigen microarray containing DENV-2 whole virus particles spotted on pGOLD slides (Nirmidas Biotech, California), as described (Robinson et al., 2019; Zhang et al., 2017). Previously defined cutoffs based on mean levels +3 standard deviations were used.

### Antibody conjugation and lyophilization

Antibody conjugation was performed as previously described (Hartmann et al., 2019). Briefly, metal-isotope labeled antibodies used in this study were conjugated using the MaxPar X8 Antibody Labeling kit per manufacturer instruction (Fluidigm) or purchased from Fluidigm pre-conjugated. Each conjugated antibody was quality checked and titrated to optimal staining concentration on healthy human PBMCs. Aliquots of the optimized surface and intracellular antibody cocktails and live cell barcoding (LCB) cocktails (Hartmann et al., 2018) were lyophilized to minimize batch effect, as previously described (Sahaf et al., 2020). Briefly, 20 uL of 500 mM trehalose in deionized, distilled water (DDW) was added to each aliquot and the final volume of each aliquot was brought to 100 uL with DDW. Aliquots were transferred uncapped into a pre-chilled tube rack in a lidded cryobox (Biocision), and placed at -80°C for one hour. Aliquots were transferred to a vacuum chamber cabinet for 24 hours. Desiccated aliquots were kept at -20°C until reconstituted for cell staining.

### Mass cytometry workflow

Samples were processed in eight batches of twenty samples. For each batch, an aliquot of a common cryopreserved PBMC sample from a healthy donor was added for batch correction. Cryopreserved PBMC samples where thawed at 37°C and transferred to 13 mL of cold cell culture medium (RPMI-1640 (Gibco), 10% FBS, 20 U/ml sodium heparin, and 0.025 U/ml benzonase (Sigma) and washed once (250g, 4°C). After cell counting, up to 1.5 million cells per sample were transferred into new tubes and washed in cell-staining media (CSM: PBS, 0.5% BSA, and 0.02% sodium azide (Sigma); 250g, 4°C). Cells were stained with LCB cocktails (reconstituted in CSM) for 30 minutes at 25°C. Cells were washed in CSM (250g, 4°C), Fc-blocked for 10 minutes (FcX, Biolegend) and stained with the surface antibody cocktail (reconstituted in CSM) for 60 minutes at 25°C. Viability staining was performed by resuspending cells in 1 mL 500 nM PdCl in PBS for 5 minutes. Cells were washed in CSM (250g, 4°C) and fixed using the FoxP3/transcription factor fixation buffer (eBioscience) for 1 hour at 25°C. Cells were washed in FoxP3/transcription factor permeabilization buffer (eBioscience) (500g, 4C) and stained with the intracellular antibody cocktail (reconstituted in permeabilization buffer) for 1 hour at 25°C. Cells were washed in permeabilization buffer (500g, 4°C) and resuspended in intercalator solution (PBS, 1.6% PFA, and 0.5 mM rhodium (Fluidigm) overnight at 4°C. The next day, cells were washed once in CSM and twice in DDW (500g, 4°C), filtered through a 35mm nylon mesh cell strainer, and resuspended in DDW with 1x EQ four-element calibration beads (Fluidigm). Barcoded samples were acquired in several fcs files over the course of a day on a CyTOF2 mass cytometer (Fluidigm) using a Super Sampler injection system (Victorian Airship).

### Data pre-processing

Fcs files from the mass cytometer were bead normalized within each batch (Finck et al., 2013) using the R package, *premessa*, except for batch 6, which lacked sufficient bead events. Files were uploaded to cellengine.com and leukocytes were gated as barium^+^ DNA^+^ bead^-^ viability^-^ and CD45^+^ (Tsai et al., 2020) (**Figure S1B**). Fcs files were downloaded and compensated with the R package, *CATALYST* (Chevrier et al., 2018). Files from batch 6 were 99^th^ percentile normalized to account for sensitivity drift over time without bead normalization. Within each batch, CD3 was quantile normalized across files to account for antibody degradation over the course of acquisition (**Figure S1C**). Files within each batch were then concatenated and debarcoded using *premessa*. Fcs files with <1000 events were removed from downstream analysis. Fcs files were transformed with hyperbolic arcsin function using a cofactor of 5.

Batch correction was performed utilizing the control sample included with each batch. For each protein channel, the median expression value was determined for all controls. The difference in medians was subtracted from all events to shift the distributions downward to have identical medians. Negative values that were introduced by this process were reset to the absolute value of a random normal distribution centered at zero to align with the noise floor for each channel. The resulting distributions were then 99^th^ percentile normalized downward. The transforms applied to each batch control sample were then applied to all samples from that batch. This approach retained biological heterogeneity while accommodating batch correction for a range of distributions and batch effect manifestations (**Figure S1D**).

### Clustering

Clustering was used to organize cells into known immune cell subsets rather than to discover novel phenotypes. In this way, expression of activation molecules, checkpoint molecules, *etc.* that were not used for clustering could be compared between clinical groups without introducing bias. We employed a supervised hierarchical clustering method that parallels manual gating, as previously described (Glass et al., 2020). Briefly, cells were over-clustered using the FlowSOM package (Van Gassen et al., 2015) and clusters were manually assigned to cell type (T, B, NK, myeloid) (**Figure 1B**, panels). For each cell type, cells were re-clustered with FlowSOM using only markers specific to that cell type (*e.g.* CD4 was used to cluster T cells, but not B cells). Clusters were then manually assigned to cell subtype (**Figure 1C**, colors) and cell population (**Figure 1B**, rows) based on known expression patterns (Alcántara-Hernández et al., 2017; Glass et al., 2020; Hartmann et al., 2019; Tsai et al., 2020).

### Feature set generation

For each sample, summary statistics were derived that were used for various comparisons. Samples with <3000 cells were removed from downstream analyses due to sampling bias. The proportions of cell populations out of total leukocytes and the proportion of cell populations out of cell subtypes were calculated. For B cells, the proportion of isotype usage and ratios of isotype usage were also calculated for each cell population. For each cell population, the mean expression of markers relevant to that cell type were quantified (*e.g.* CTLA-4 was quantified on T cells but not myeloid cells). Mean expression was chosen as changes in the mean are sensitive to both a shift in protein copy number as well as a shift in the percent of positive cells. Percent positive was not used as a feature due to high correlation with the mean.

### Dimensionality reduction

The LDA (**Figure 1E**) was generated with the *MASS* package using all features significantly different between any pairwise comparison of acute clinical status samples. The manifold was trained to separate samples by clinical status. The UMAP (**Figure 1C**) was generated using the *uwot* package with min_dist=0.7 and n_neighbors=15 with all proteins as input and an equal subsampling of patients and clinical statuses. UMAP Coordinates were initialized with LDA coordinates, trained to separate cell subtype (Amouzgar et al., 2022).

### Statistical analyses

All pairwise comparisons between samples employed the Wilcoxon rank sum test for all features. For comparisons between acute samples, multiple hypothesis correction was performed using FDR. Features were only considered significantly different if q<0.05 and |effect|>0.5. For comparisons between convalescent samples, timepoints, children, adults, and SD subtypes, multiple hypothesis correction was not used due to small sample sizes. To maintain statistical rigor, the effect size requirement was increased, so features were only considered significant if p<0.05 and |effect|>1.0). For correlation analysis, the Pearson method was employed and with multiple hypothesis correction using FDR when multiple correlations were calculated.

## Code and data availability

Fcs files will be made available at flowrepository.com. Code to generate all figures available at github.com/davidrglass/dengue.

## Author Contributions

Conceptualization, S.E, M.L.R., V.D.; Methodology, D.R.G.*, M.L.R*, S.C.B., M.K.S., B.A.P., H.M., S.E.; Software, D.R.G, M.L.R; Formal Analysis, D.R.G., M.L.R.; Investigation, M.L.R*, D.R.G.*, V.D., F.J.H., M.B.; Resources, S.E., S.C.B, O.L.A.R., A.N.S., M.C., R.M.G., N.B., J.G.M., M.I.E.C, L.A,V.C, E.M.R.G., F.R.; Data Curation, M.L.R., D.R.G, O.L.A.R., A.N.S., M.C., R.M.G., N.B., M.I.E.C, L.A,V.C, E.M.R.G., F.R. Writing – Original Draft, M.L.R.*, D.R.G.*, S.E; Writing – Review and Editing, M.L.R., D.R.G., S.E., V.D.,S.C.B.; Visualization, D.R.G, M.L.R., V.D.; Supervision, S.E., S.C.B.; Funding Acquisition, S.E, S.C.B. *denotes equal contribution.

## Acknowledgements

This work was supported by Catalyst and Transformational Awards from Dr. Ralph & Marian Falk Medical Research Trust, an Investigator Initiated Award number W81XWH1910235 from the Department of Defense (DoD) office of the Congressionally Directed Medical Research Programs (CDMRP)/Peer Reviewed Medical Research Program (PRMRP), and a National Institute of Allergy and Infectious Diseases (NIAID) grant U19 AI057229 to S.E.. S.E. is a Chan Zuckerberg Biohub investigator, who is also supported by an Investigator Initiated Award number W81XWH2210283 and an expansion award number W81XWH2110456 from the DoD office of the CDMRP/PRMRP, a NIAID grant RO1AI158569, and a Defense Threat Reduction Fundamental Research to Counter Weapons of Mass Destruction grant HDTRA11810039. M.L.R. was supported by the A.P. Giannini Foundation Postdoctoral Fellowship and the Harold Amos Medical Faculty Development Program. D.R.G. was supported by a Stanford Graduate Fellowship and a Bio-X Stanford Interdisciplinary Graduate Fellowship. V.D. was supported by a Chan Zuckerberg Biohub Collaborative Postdoctoral Fellowship. F.J.H. was supported by an EMBO Long-Term Fellowship ALTF 1141–2017, the Novartis Foundation for Medical-Biological Research 16C148, and the Swiss National Science Foundation SNF Early Postdoc Mobility P2ZHP3-171741. S.C.B. was supported by NIH awards DP2EB024246, U24CA224309, R01AG057915, R01AG056287, R01AG068279, 1UH3CA246633.

