## Supplemental Figures for "Magnitude and kinetics of the human immune cell response associated with severe dengue progression by single-cell proteomics"

**Supplemental text:**

#### Challenges in identifying DENV-infected cells in patient-derived PBMCs via mass cytometry

The precise immune cell targets of DENV in the human blood remain incompletely characterized. Myeloid cell types including monocytes and dendritic cells have been shown to be actively infected with DENV (Durbin et al., 2008; Kyle et al., 2007). However a previous study has shown that B cells can maintain DENV infection (Upasani et al., 2021) as well, and we have demonstrated that B cell populations, specifically naïve B cells, harbor the large bulk of viral RNA (Zanini et al., 2018). Here, we aimed to further define DENV target cells by conjugating three antibodies targeting the DENV envelope (structural), and NS1 and NS3 (nonstructural) proteins (NS1, NS3). Titration of these anti-DENV antibodies in DENV infected Huh7 cells revealed clear bimodal signal and an inoculum dependent increase in DENV protein marker staining at 48 hours postinfection **(Figure S4B).** However, when used to detect DENV in primary patient-derived PBMCs, we detected a greatly increased noise floor, resulting in a small fraction of cells that stained positive for all three DENV proteins **(Figure S4C).** Moreover, there was no correlation between the expression level of the DENV proteins and DENV RNA level measured via RT-qPCR in patient serum **(Figure S4D).** We speculate that the DENV-specific antibodies bound viral protein debris non-specifically associated with immune cell membranes during cell staining, preventing discrimination of infected cells. Further optimization will therefore be required to detect primary DENV-infected cells in PBMCs via CyTOF.

### Figure S1

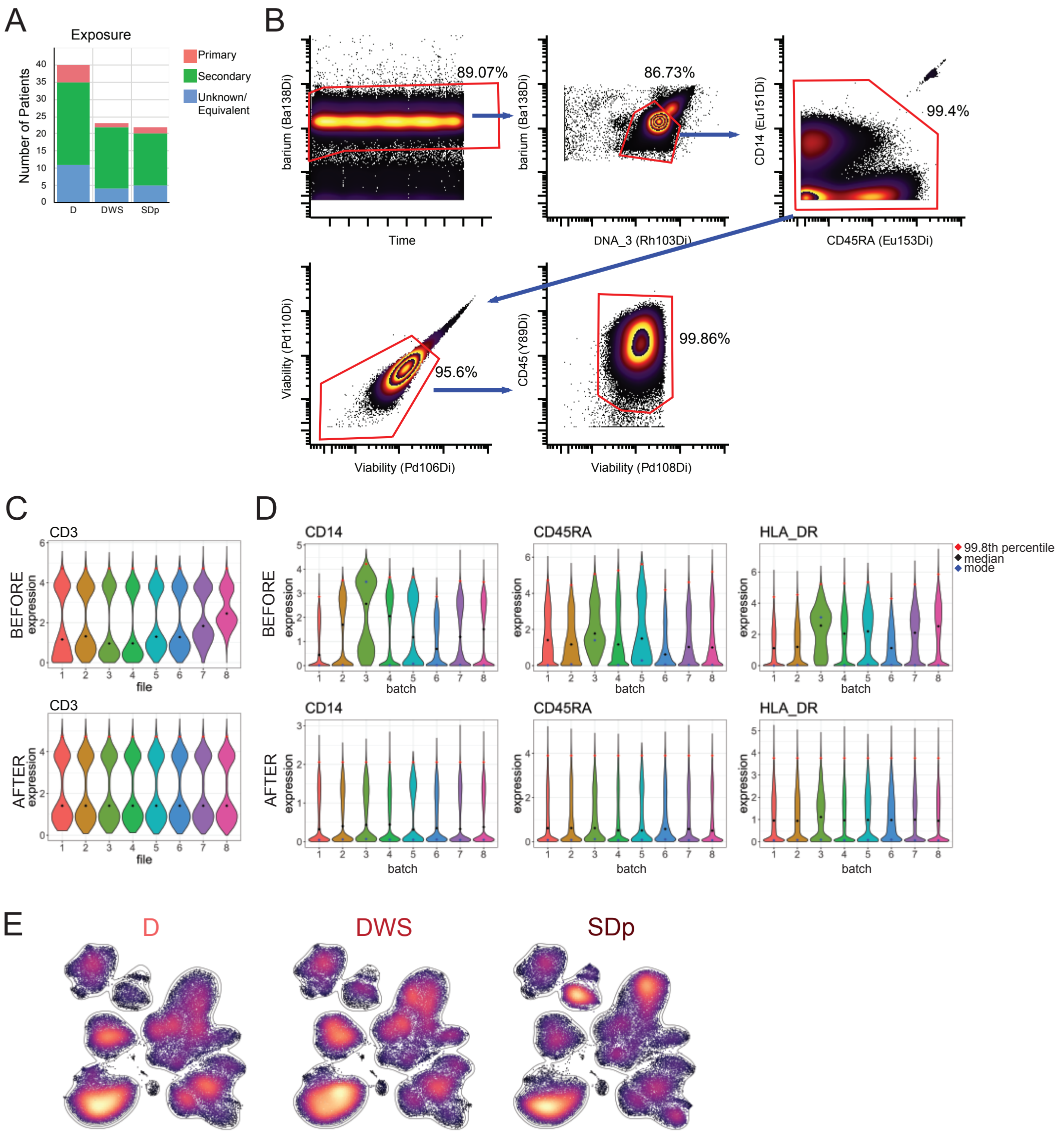


**Figure S1: Patient information, gating, normalization, and unbiased segregation of cells by clinical status – related to Figure 1**

A) Composition of cohort by dengue exposure status (colors) and clinical status (columns).

B) Exemplative leukocyte gating strategy

C) Exemplative correction of CD3 within a batch. Violins represent cell distributions of barcoded files not yet deconvolved that were iteratively collected over the course of a day.

D) Exemplative batch correction of three protein channels. Violins represent cell distributions from a common control sample run with each batch.

E) UMAP plots with same coordinates as in Figure 1C, separated by disease status and colored by density.

### Figure S2

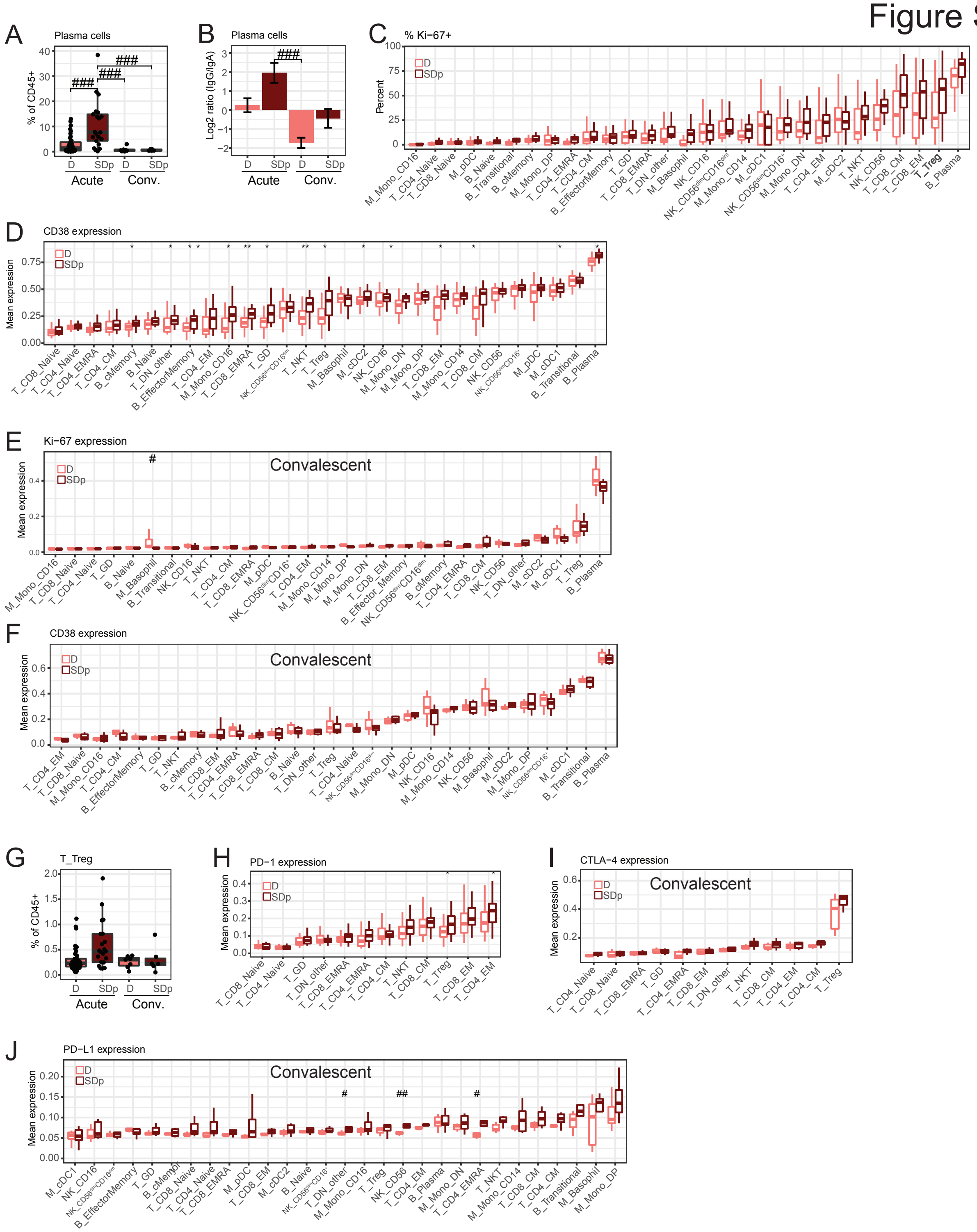


**Figure S2. Additional features of immune activation and regulation in acute and convalescent SD and D samples – related to Figure 2**

(A, G) Plasma cell and Treg abundance by clinical status and time. Dots represent patients.

(B) Mean $log_2$ ratio of patient $IgG^+$ plasma cell abundance to $IgA^+$ plasma cell abundance by clinical status and time. Error bars represent standard error of mean (SEM).

(C) % Ki-67+ by cell population and clinical status in acute samples.

(D, H) Mean CD38 (D) and PD-1 (H) expression by cell population and clinical status in acute samples.

(E, F, I, J) Mean Ki-67, CD38, CTLA-4, and PD-L1 expression by cell population and clinical status in convalescent samples.

In boxplots, center line signifies median, box signifies interquartile range (IQR) and whiskers signify IQR +/- 1.5*IQR. q<0.05 & |effect|>0.5; ** q<0.01 & |effect|>0.5; *** q<0.005 & |effect|>0.5; # p<0.05 & |effect|>1.0; # p<0.05 & |effect|>1.0; ## p<0.01 & |effect|>1.0; ### p<0.005 & |effect|>1.0 by Wilcoxon rank sum tests. D, dengue; SDp, SD progressors.

Figure S3

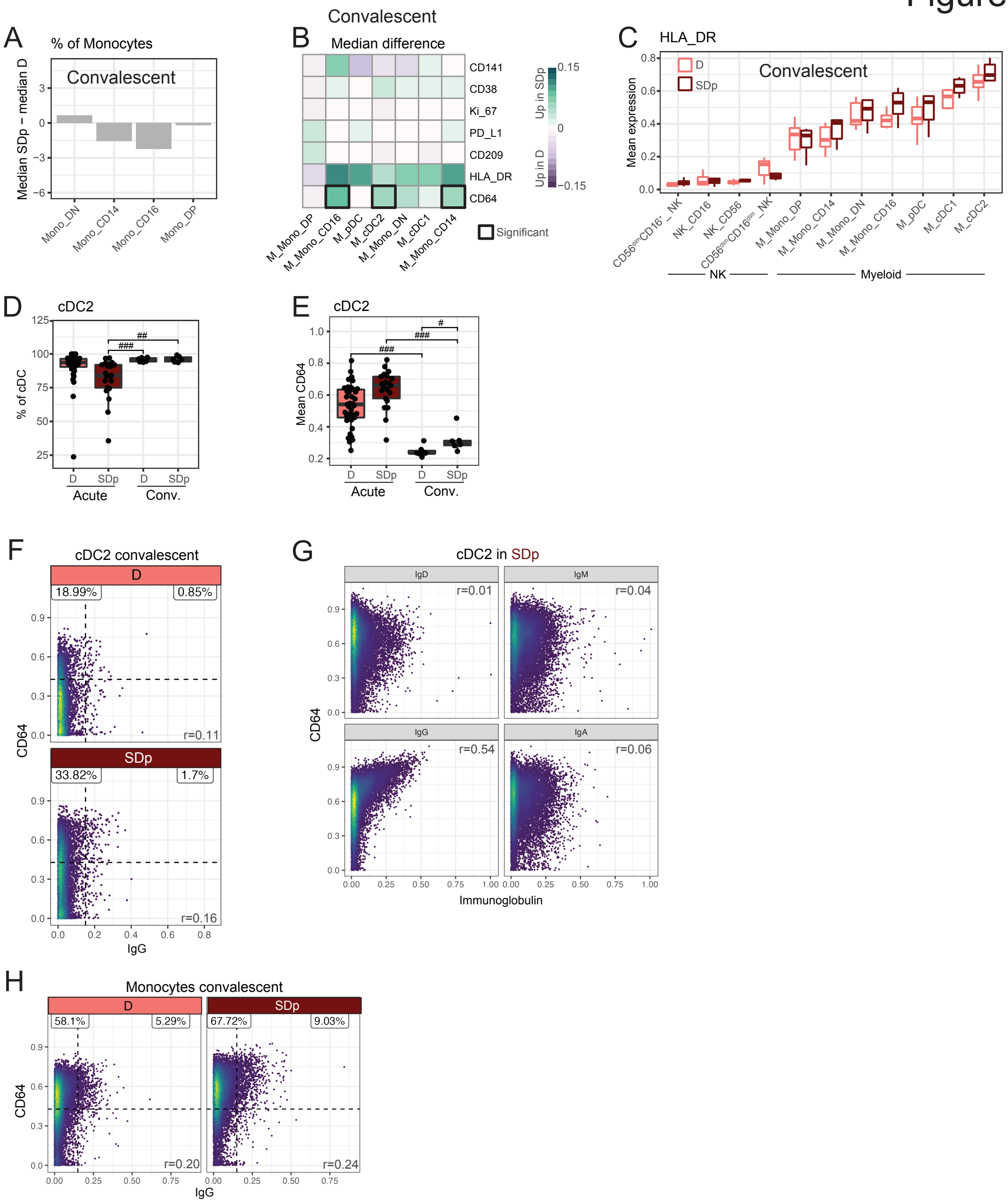


**Figure S3: Additional innate immune features of acute and convalescent SDp and D samples – related to Figure 3**

(A) Difference in $\log_2$ ratio of median relative abundances of monocyte subtypes (columns) between convalescent SDp and D patients out of total monocytes. Teal bars indicate significance (q<0.05 & |effect|>0.5).

(B) Difference in cohort median of patient mean expression between convalescent SDp and D samples across molecules (rows) in myeloid cell populations (columns). Black boxes indicate significance ($p<0.05$ & |effect|>0.5).
(C) Mean HLA-DR expression in NK and myeloid cell subtypes by clinical status in convalescent samples.
(D, E) Box plots of cDC2 fraction of CD45+ cells (D) and mean CD64 expression (E) by clinical status in acute and convalescent samples. Dots represent individual patients.
(F, H) Scatter plots of single cell CD64 expression versus IgG detection in convalescent samples in cDC2s (F) and monocytes (H) by clinical status. Percent proportion of cells in each quadrant, derived from an equal subsampling of cells by clinical status and patient are shown.
(G) Scatter plots of single cell CD64 expression versus immunoglobulin isotypes detection on cDC2s in acute SDp samples. Cells were derived from an equal subsampling of cDC2s by patient.
In boxplots, center line signifies median, box signifies interquartile range (IQR) and whiskers signify IQR +/- 1.5*IQR. Dots in A, C, D, F, and H represent individual patients.
* $q<0.05$ & |effect|>0.5; ** $q<0.01$ & |effect|>0.5; *** $q<0.005$ & |effect|>0.5; # $p<0.05$ & |effect|>1.0; ## $p<0.01$ & |effect|>1.0; ### $p<0.005$ & |effect|>1.0 by Wilcoxon rank sum tests. Q-values represent FDR-corrected p-values. r in F, G, H and I was calculated by Pearson correlation. D, dengue; SDp, SD progressors; conv., convalescent.

### Figure S4

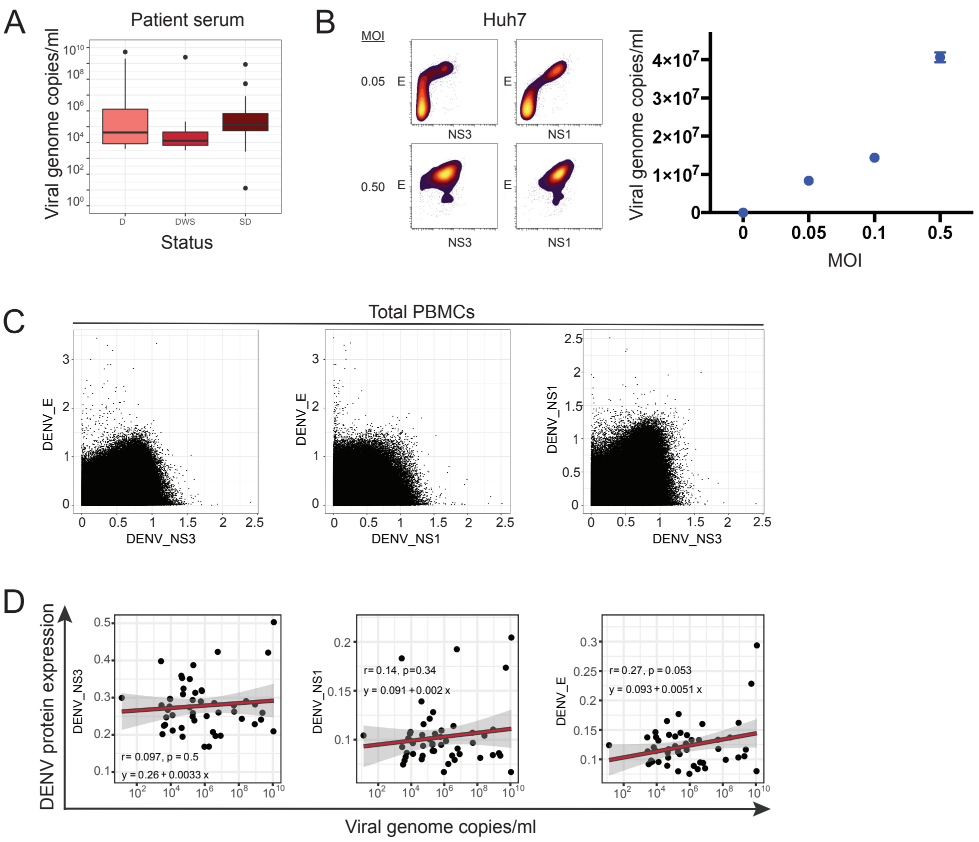


**Figure S4: Quantification of DENV viral load and evaluation of DENV target cells via CyTOF**

(A) Boxplot depicting level of viremia determined by qRT-PCR by dengue clinical status. In boxplots, center line signifies median, box signifies interquartile range (IQR) and whiskers signify IQR +/- 1.5*IQR.

(B) CyTOF analysis of Huh7 cells infected with DENV-2 at MOIs of 0.05 and 0.5. Left panel: Dot plots of mean signal intensity of DENV E, NS1 and NS3 proteins. Right panel: Viral load measured via qRT-PCR for various MOIs.

(C) Contour plots of single cell DENV E, NS1 and NS3 protein expression in total PBMCs derived from the dengue Colombia cohort.

(D) Scatter plots of mean DENV E, NS1 and NS3 protein expression in PBMCs versus viral load in serum samples with detectable virus by qRT-PCR. Fitting line and 95% CI (shading) were derived by linear regression. Correlation coefficient (r) and p-values were calculated by Pearson correlation.

E, Envelope; NS1, nonstructural 1; NS3, nonstructural 3.

Figure S5

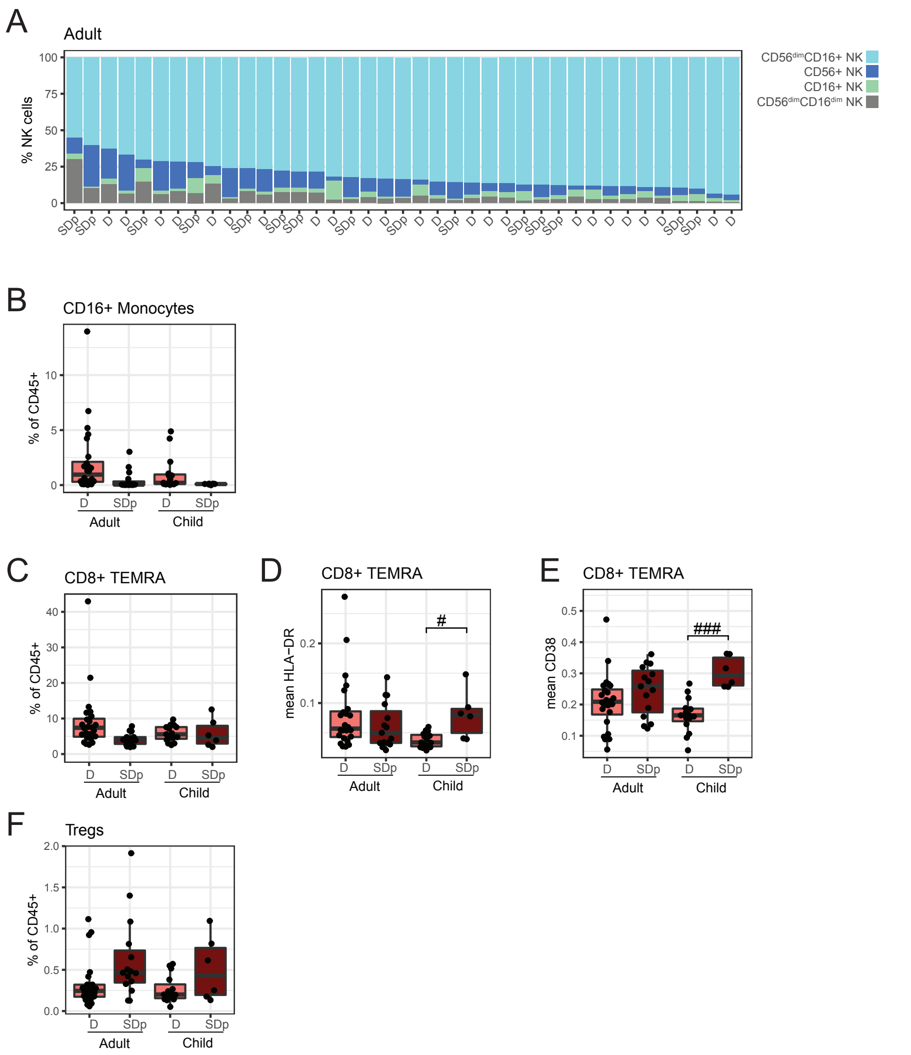


**Figure S5: Additional features differentiating SDp children from SDp adults – related to Figure 4**

(A) Proportion of NK cell subtypes in adults ordered by $CD56^{dim}CD16^{+}$ NK cell abundance. Columns represent individual patients, labeled by clinical status.

(B, C, F) Box plots of $CD16^{+}$ monocytes (B) $CD8^{+}$ TEMRA (C), Treg (F)

(F) fractions of CD45+ cells by clinical status and age. Dots represent individual patients.

(D, E) Box plots of mean HLA-DR (C) and CD38 (D) expression in CD8+ TEMRA cells by clinical status and age. Dots represent patients.

In boxplots, center line signifies median, box signifies interquartile range (IQR) and whiskers signify IQR +/- 1.5*IQR. # $p<0.05$ & |effect|>1.0; ## $p<0.01$ & |effect|>1.0; ### $p<0.005$ & |effect|>1.0 by Wilcoxon rank sum tests. D, dengue; SDp, SD progressors.

Figure S6

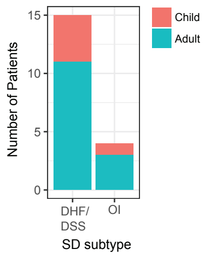


**Figure S6: SDp patient distribution by SD subtypes and age – related to Figure 5**

Patient composition of SD samples by age (colors) and SD subtype (columns).

DHF/DSS, Dengue hemorrhagic fever/dengue shock syndrome; OI, organ impairment.

Figure S7

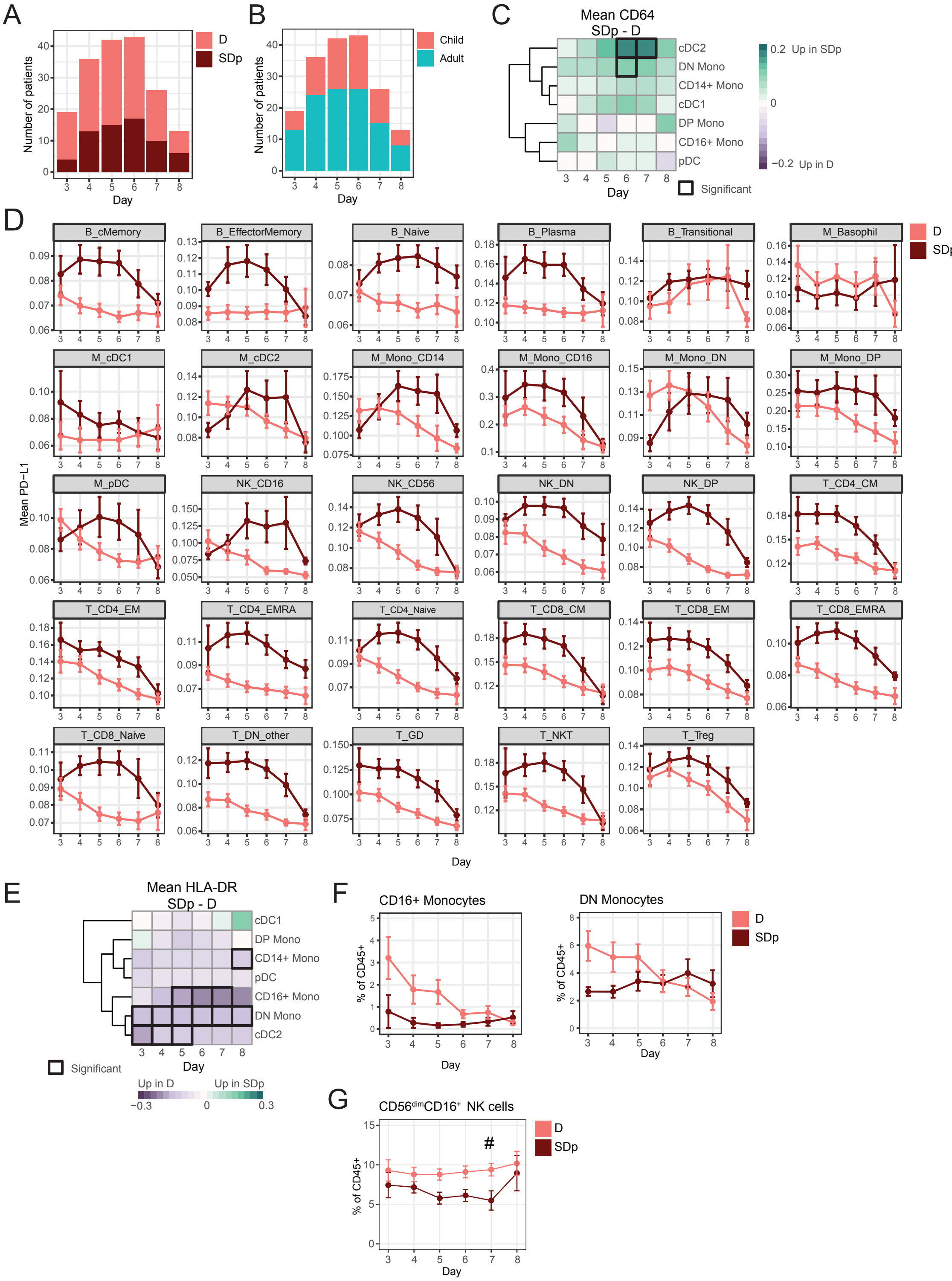
A
Number of patients
Day
D
SDp
B
Number of patients
Day
Child
Adult
C
Mean CD64
SDp - D
cDC2
DN Mono
CD14+ Mono
cDC1
DP Mono
CD16+ Mono
pDC
Day
0.2 Up in SDp
0
−0.2 Up in D
Significant
D
Mean PD−L1
B_cMemory
B_EffectorMemory
B_Naive
B_Plasma
B_Transitional
M_Basophil
M_cDC1
M_cDC2
M_Mono_CD14
M_Mono_CD16
M_Mono_DN
M_Mono_DP
M_pDC
NK_CD16
NK_CD56
NK_DN
NK_DP
T_CD4_CM
T_CD4_EM
T_CD4_EMRA
T_CD4_Naive
T_CD8_CM
T_CD8_EM
T_CD8_EMRA
T_CD8_Naive
T_DN_other
T_GD
T_NKT
T_Treg
Day
D
SDp
E
Mean HLA-DR
SDp - D
cDC1
DP Mono
CD14+ Mono
pDC
CD16+ Mono
DN Mono
cDC2
Day
Significant
Up in D
Up in SDp
−0.3
0
0.3
F
CD16+ Monocytes
DN Monocytes
% of CD45+
Day
D
SDp
G
CD56dimCD16+ NK cells
% of CD45+
#
Day
D
SDp

**Figure S7: Additional features of acute infection by time – related to Figure 6**

(A) Composition of clinical status (colors) by day.

(B) Composition of age (colors) by day.

(C, E) Difference in cohort mean of patient mean CD64 (D) and HLA-DR (E) expression between SDp and D across cell populations (rows) by day (columns). Black boxes indicate significance ($p<0.05$ & $|effect|>1.0$).

(D) Cohort mean of patient mean PD-L1 expression by day (columns) and clinical status (color), in the indicated cell populations. Each graph is individually scaled.

(F) Cohort mean of $CD16^{+}$ (left) and DN (right) monocyte abundances of CD45+ cells by day (columns) and clinical status (color).

(G) Cohort mean of $CD56^{dim}CD16^{+}$ NK cell abundance of CD45+ cells by day (columns) and clinical status (color).

Error bars represent SEM. D, dengue; SDp, SD progressor; DN, double negative.

Figure S1

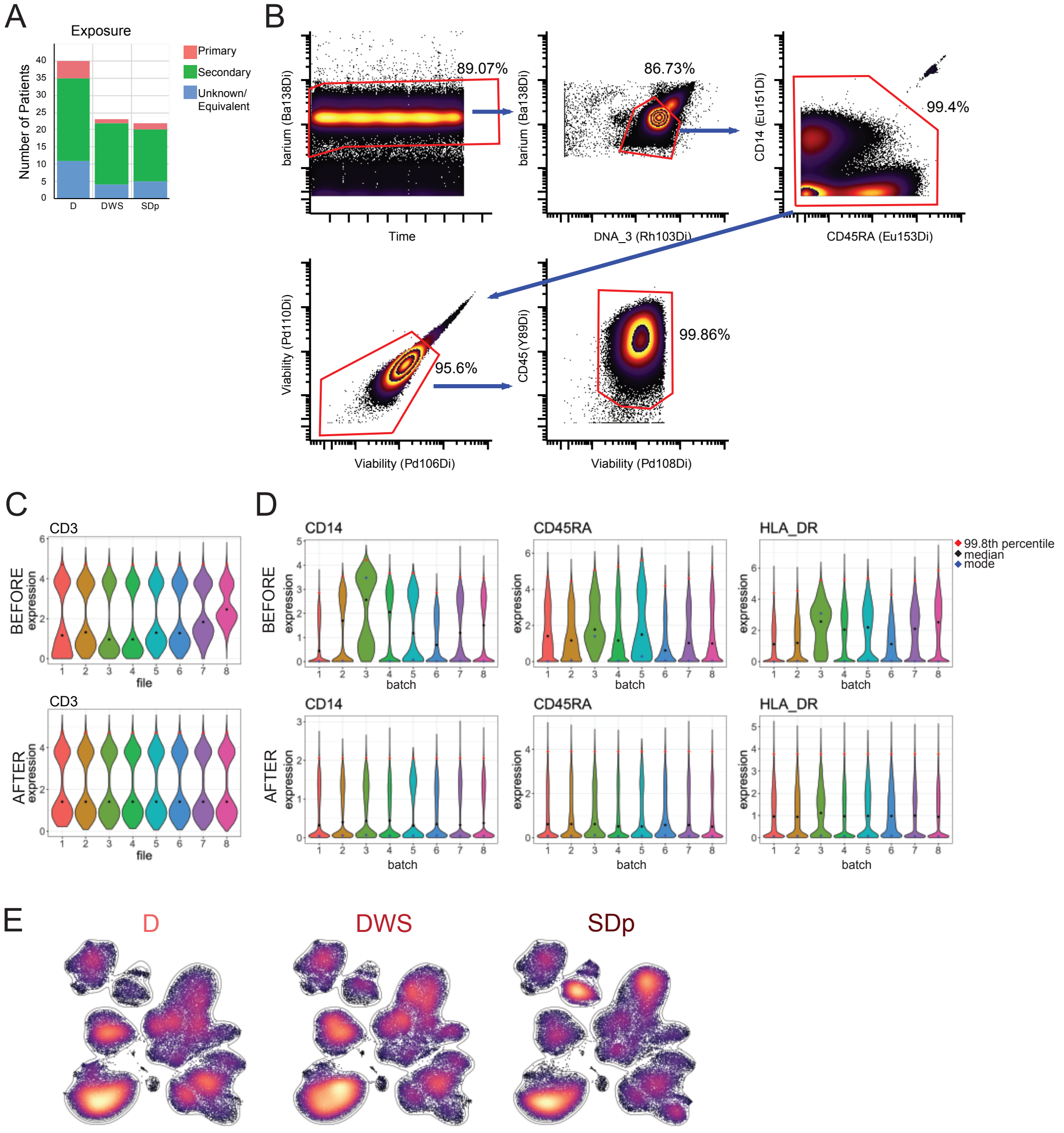


**Figure S1: Patient information, gating, normalization, and unbiased segregation of cells by clinical status – related to Figure 1**

A) Composition of cohort by dengue exposure status (colors) and clinical status (columns).

B) Exemplative leukocyte gating strategy

C) Exemplative correction of CD3 within a batch. Violins represent cell distributions of barcoded files not yet deconvolved that were iteratively collected over the course of a day.

D) Exemplative batch correction of three protein channels. Violins represent cell distributions from a common control sample run with each batch.

E) UMAP plots with same coordinates as in Figure 1C, separated by disease status and colored by density.

Figure S2

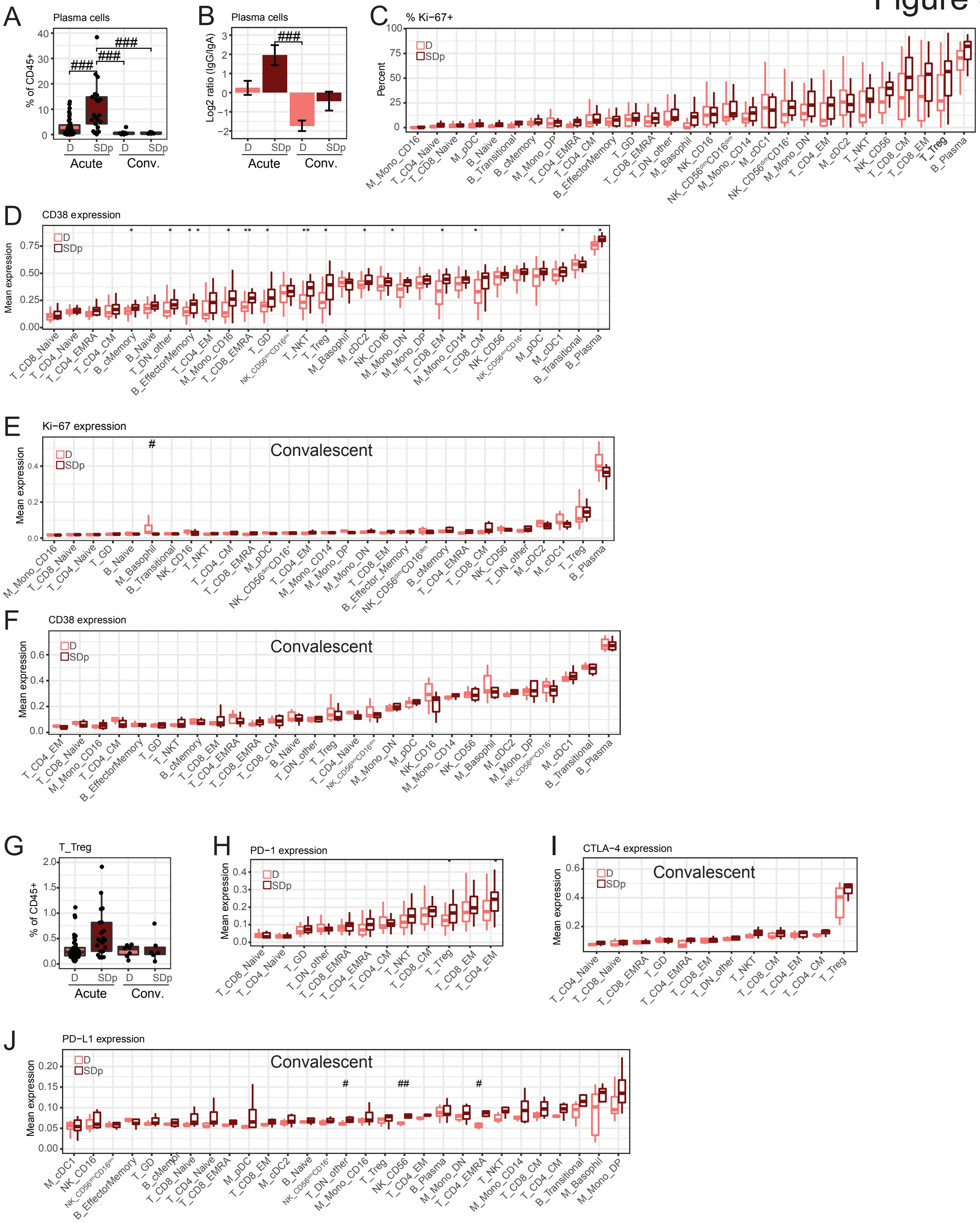


**Figure S2. Additional features of immune activation and regulation in acute and convalescent SD and D samples – related to Figure 2**

(A, G) Plasma cell and Treg abundance by clinical status and time. Dots represent patients.
(B) Mean $log_2$ ratio of patient $IgG^+$ plasma cell abundance to $IgA^+$ plasma cell abundance by clinical status and time. Error bars represent standard error of mean (SEM).
(C) % Ki-67+ by cell population and clinical status in acute samples.
(D, H) Mean CD38 (D) and PD-1 (H) expression by cell population and clinical status in acute samples.
(E, F, I, J) Mean Ki-67, CD38, CTLA-4, and PD-L1 expression by cell population and clinical status in convalescent samples.
In boxplots, center line signifies median, box signifies interquartile range (IQR) and whiskers signify IQR +/- 1.5*IQR. q<0.05 & |effect|>0.5; ** q<0.01 & |effect|>0.5; *** q<0.005 & |effect|>0.5; # p<0.05 & |effect|>1.0; # p<0.05 & |effect|>1.0; ## p<0.01 & |effect|>1.0; ### p<0.005 & |effect|>1.0 by Wilcoxon rank sum tests. D, dengue; SDp, SD progressors.

Figure S3

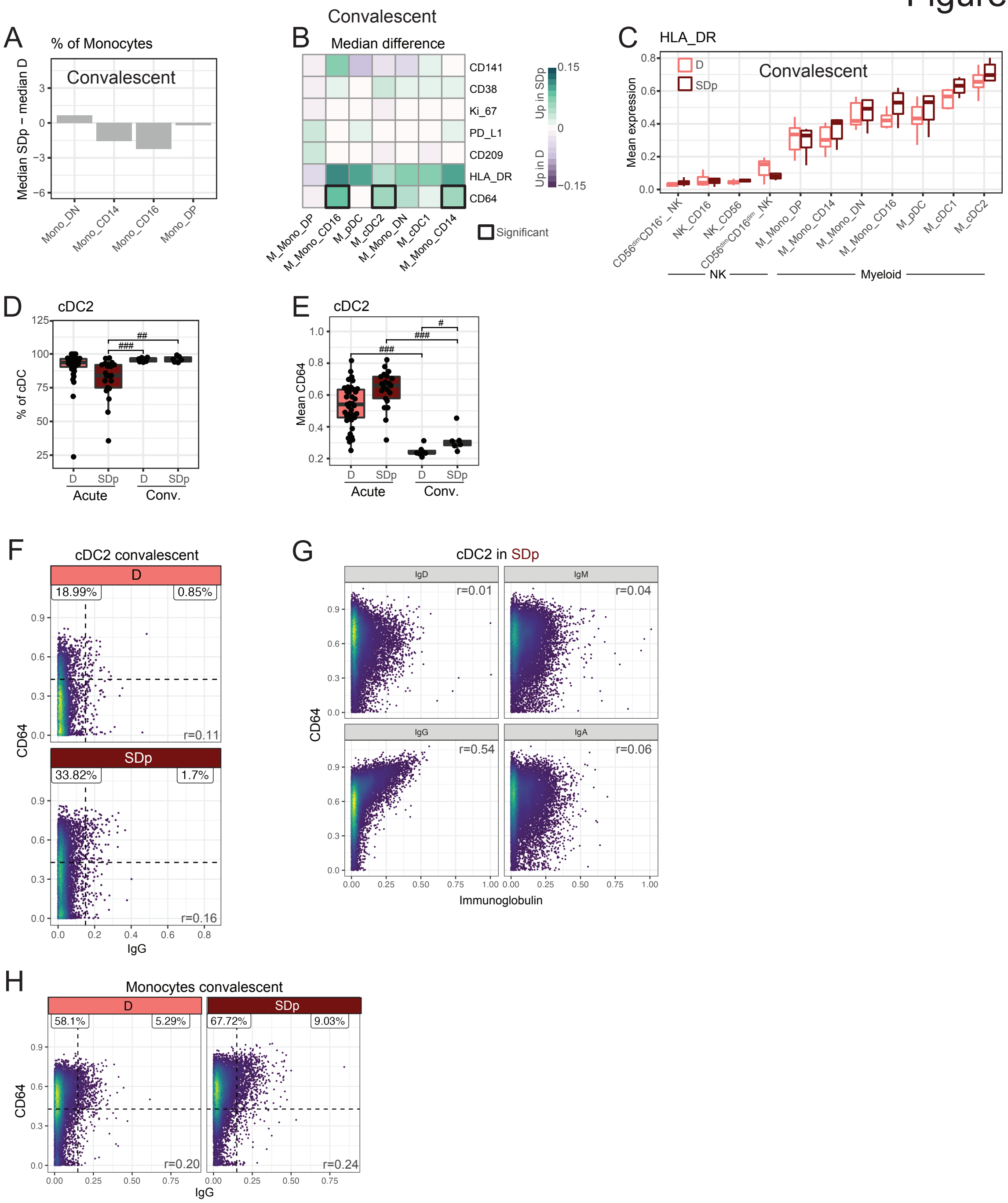


**Figure S3: Additional innate immune features of acute and convalescent SDp and D samples – related to Figure 3**

(A) Difference in $\log_2$ ratio of median relative abundances of monocyte subtypes (columns) between convalescent SDp and D patients out of total monocytes. Teal bars indicate significance (q<0.05 & |effect|>0.5).

(B) Difference in cohort median of patient mean expression between convalescent SDp and D samples across molecules (rows) in myeloid cell populations (columns). Black boxes indicate significance ($p<0.05$ & |effect|>0.5).
(C) Mean HLA-DR expression in NK and myeloid cell subtypes by clinical status in convalescent samples.
(D, E) Box plots of cDC2 fraction of CD45+ cells (D) and mean CD64 expression (E) by clinical status in acute and convalescent samples. Dots represent individual patients.
(F, H) Scatter plots of single cell CD64 expression versus IgG detection in convalescent samples in cDC2s (F) and monocytes (H) by clinical status. Percent proportion of cells in each quadrant, derived from an equal subsampling of cells by clinical status and patient are shown.
(G) Scatter plots of single cell CD64 expression versus immunoglobulin isotypes detection on cDC2s in acute SDp samples. Cells were derived from an equal subsampling of cDC2s by patient.
In boxplots, center line signifies median, box signifies interquartile range (IQR) and whiskers signify IQR +/- 1.5*IQR. Dots in A, C, D, F, and H represent individual patients.
* $q<0.05$ & |effect|>0.5; ** $q<0.01$ & |effect|>0.5; *** $q<0.005$ & |effect|>0.5; # $p<0.05$ & |effect|>1.0; ## $p<0.01$ & |effect|>1.0; ### $p<0.005$ & |effect|>1.0 by Wilcoxon rank sum tests. Q-values represent FDR-corrected p-values. r in F, G, H and I was calculated by Pearson correlation. D, dengue; SDp, SD progressors; conv., convalescent.

Figure S4

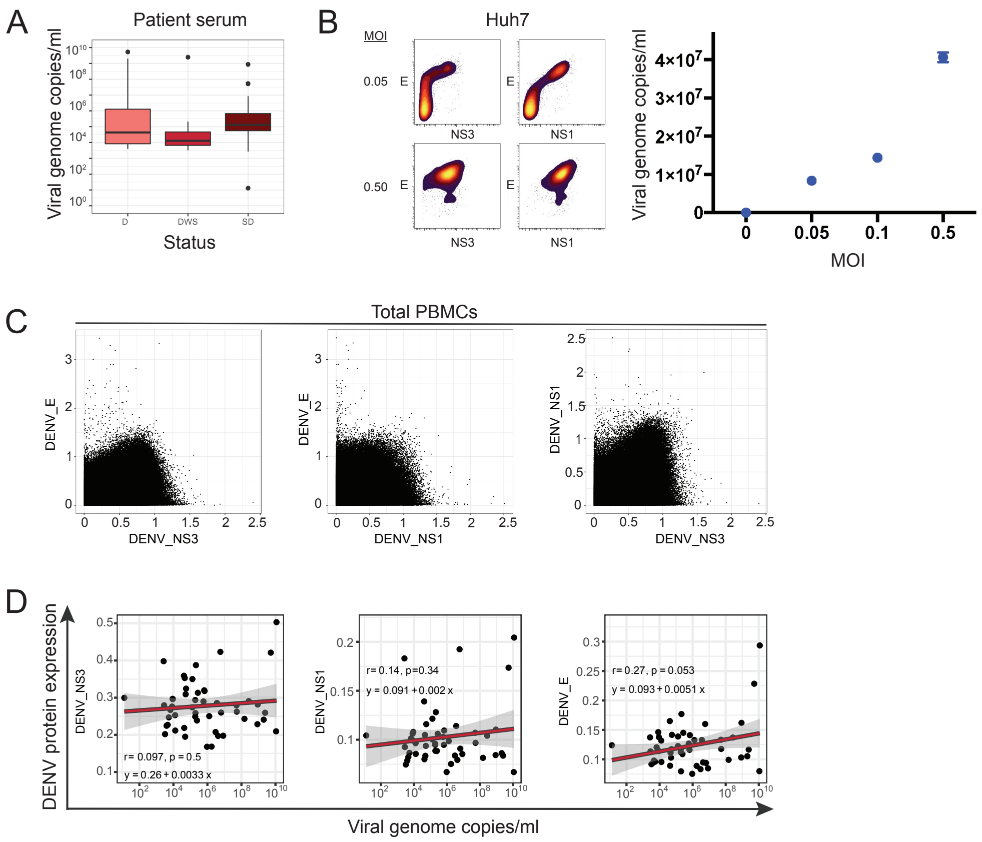


**Figure S4: Quantification of DENV viral load and evaluation of DENV target cells via CyTOF**

(A) Boxplot depicting level of viremia determined by qRT-PCR by dengue clinical status. In boxplots, center line signifies median, box signifies interquartile range (IQR) and whiskers signify IQR +/- 1.5*IQR.

(B) CyTOF analysis of Huh7 cells infected with DENV-2 at MOIs of 0.05 and 0.5. Left panel: Dot plots of mean signal intensity of DENV E, NS1 and NS3 proteins. Right panel: Viral load measured via qRT-PCR for various MOIs.

(C) Contour plots of single cell DENV E, NS1 and NS3 protein expression in total PBMCs derived from the dengue Colombia cohort.

(D) Scatter plots of mean DENV E, NS1 and NS3 protein expression in PBMCs versus viral load in serum samples with detectable virus by qRT-PCR. Fitting line and 95% CI (shading) were derived by linear regression. Correlation coefficient (r) and p-values were calculated by Pearson correlation.

E, Envelope; NS1, nonstructural 1; NS3, nonstructural 3.

### Figure S5

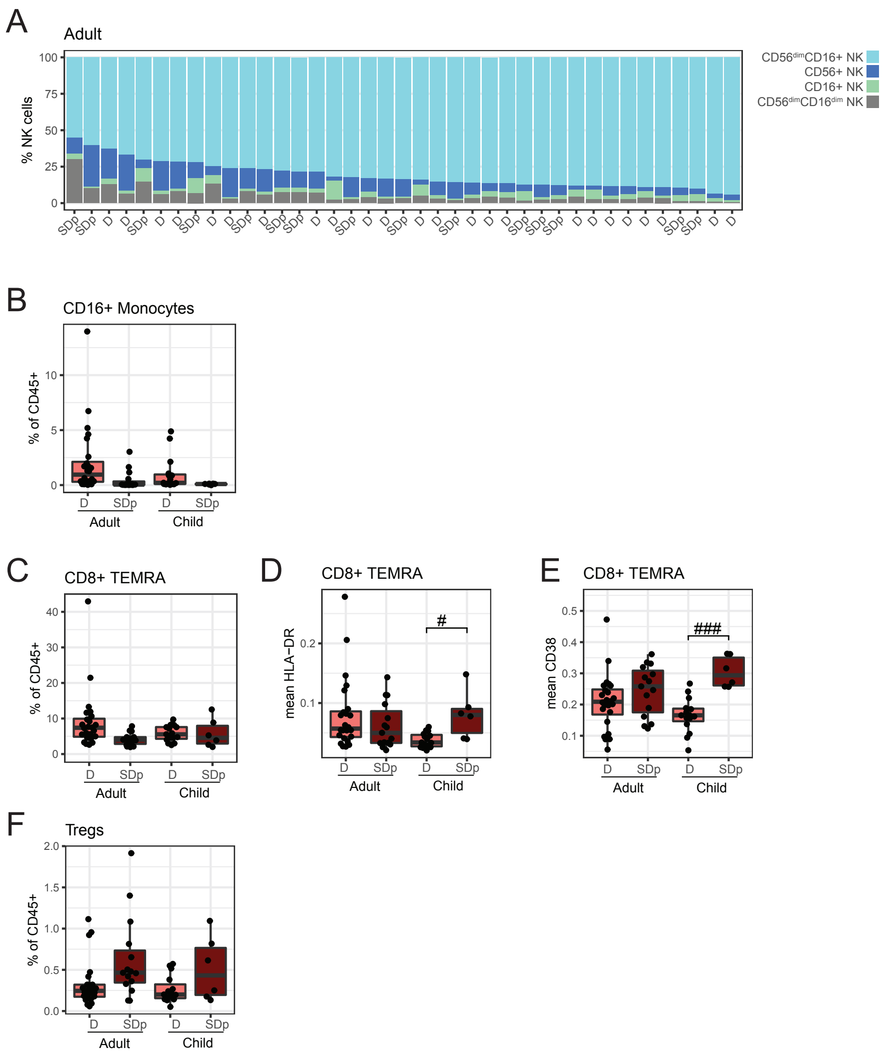


**Figure S5: Additional features differentiating SDp children from SDp adults – related to Figure 4**

(A) Proportion of NK cell subtypes in adults ordered by CD56$^{dim}$CD16$^{+}$ NK cell abundance. Columns represent individual patients, labeled by clinical status.

(B, C, F) Box plots of CD16$^{+}$ monocytes (B) CD8$^{+}$ TEMRA (C), Treg (F)

(F) fractions of CD45+ cells by clinical status and age. Dots represent individual patients.

(D, E) Box plots of mean HLA-DR (C) and CD38 (D) expression in CD8+ TEMRA cells by clinical status and age. Dots represent patients.

In boxplots, center line signifies median, box signifies interquartile range (IQR) and whiskers signify IQR +/- 1.5*IQR. # p<0.05 & |effect|>1.0; ## p<0.01 & |effect|>1.0; ### p<0.005 & |effect|>1.0 by Wilcoxon rank sum tests. D, dengue; SDp, SD progressors.

Figure S6

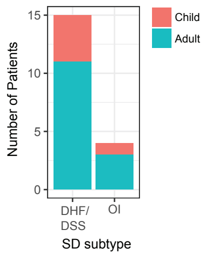


**Figure S6: SDp patient distribution by SD subtypes and age – related to Figure 5**

Patient composition of SD samples by age (colors) and SD subtype (columns).

DHF/DSS, Dengue hemorrhagic fever/dengue shock syndrome; OI, organ impairment.

Figure S7

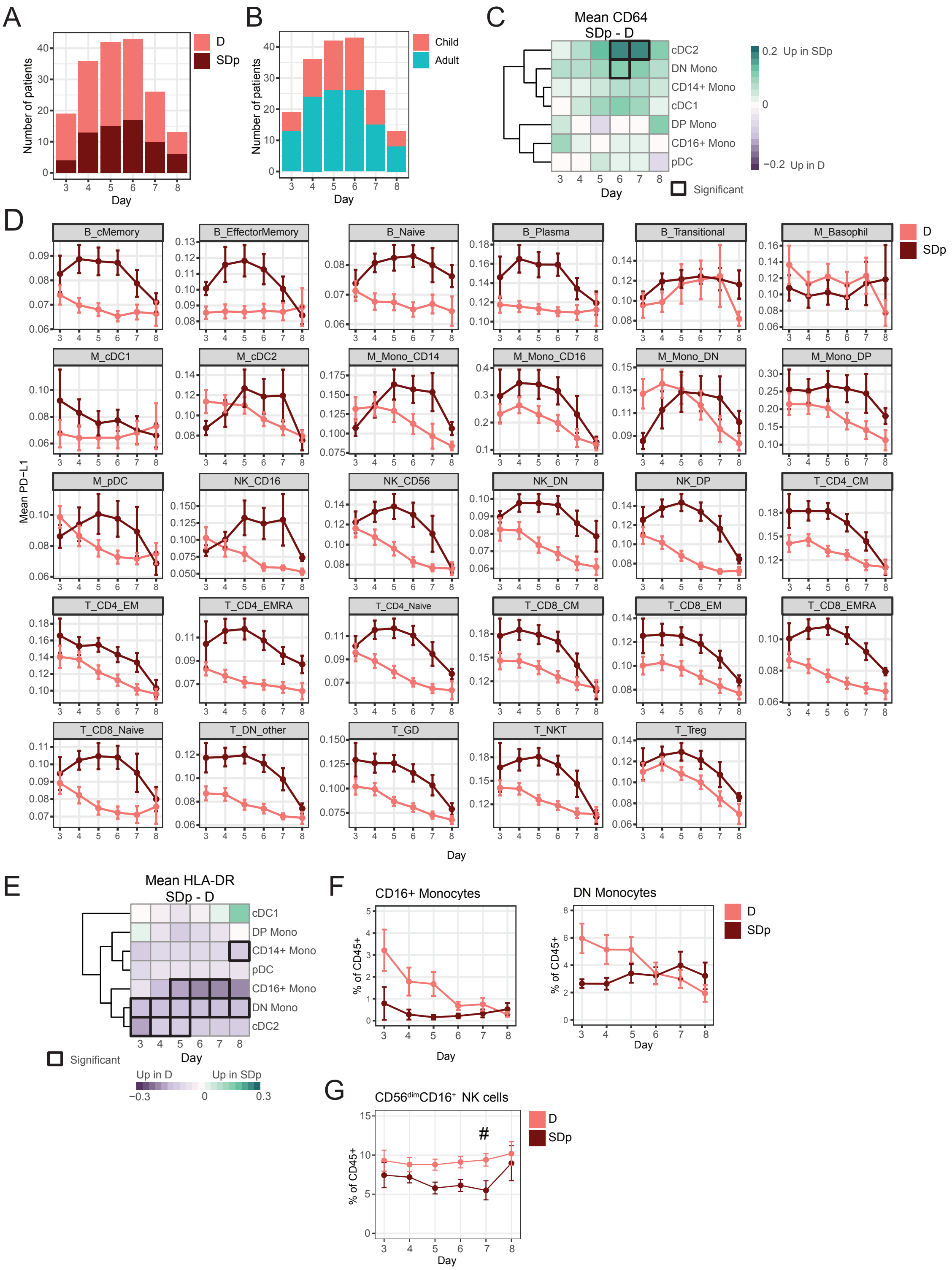
A
Number of patients
Day
D
SDp
B
Number of patients
Day
Child
Adult
C
Mean CD64
SDp - D
cDC2
DN Mono
CD14+ Mono
cDC1
DP Mono
CD16+ Mono
pDC
Day
0.2 Up in SDp
0
−0.2 Up in D
Significant
D
Mean PD−L1
B_cMemory
B_EffectorMemory
B_Naive
B_Plasma
B_Transitional
M_Basophil
M_cDC1
M_cDC2
M_Mono_CD14
M_Mono_CD16
M_Mono_DN
M_Mono_DP
M_pDC
NK_CD16
NK_CD56
NK_DN
NK_DP
T_CD4_CM
T_CD4_EM
T_CD4_EMRA
T_CD4_Naive
T_CD8_CM
T_CD8_EM
T_CD8_EMRA
T_CD8_Naive
T_DN_other
T_GD
T_NKT
T_Treg
Day
D
SDp
E
Mean HLA-DR
SDp - D
cDC1
DP Mono
CD14+ Mono
pDC
CD16+ Mono
DN Mono
cDC2
Day
Significant
Up in D
Up in SDp
−0.3
0
0.3
F
CD16+ Monocytes
DN Monocytes
% of CD45+
Day
D
SDp
G
CD56dimCD16+ NK cells
% of CD45+
#
Day
D
SDp

**Figure S7: Additional features of acute infection by time – related to Figure 6**

(A) Composition of clinical status (colors) by day.

(B) Composition of age (colors) by day.

(C, E) Difference in cohort mean of patient mean CD64 (D) and HLA-DR (E) expression between SDp and D across cell populations (rows) by day (columns). Black boxes indicate significance ($p<0.05$ & $|effect|>1.0$).

(D) Cohort mean of patient mean PD-L1 expression by day (columns) and clinical status (color), in the indicated cell populations. Each graph is individually scaled.

(F) Cohort mean of $CD16^+$ (left) and DN (right) monocyte abundances of CD45+ cells by day (columns) and clinical status (color).

(G) Cohort mean of $CD56^{dim}CD16^+$ NK cell abundance of CD45+ cells by day (columns) and clinical status (color).

Error bars represent SEM. D, dengue; SDp, SD progressor; DN, double negative.
